# Consistent alterations of human fecal microbes after transplanted to germ-free mice

**DOI:** 10.1101/495663

**Authors:** Yanze Li, Wenming Cao, Na L Gao, Xing-Ming Zhao, Wei-Hua Chen

**Author notes:** Correspondence should be addressed to Wei-Hua Chen or Xing-Ming Zhao.

## Abstract

**Background:** Fecal microbiota transplant (FMT) of human fecal samples to germ-free (GF) mice is useful for establishing causal relationships between gut microbiota and human phenotypes. However, due to intrinsic differences between human and mouse intestines and distinct diets between the two organisms, replicating human phenotypes in mouse through FMT is not guaranteed; similarly, treatments that are effective in mouse models do not guarantee their success in human either.

**Results:** In this study, we aimed to identify human gut microbes that have undergone significant and consistent changes after transplanted to GF mice across multiple experimental settings. By comparing gut microbiota profiles in 1,713 human-mouse pairs, we found strikingly on average <50% of the human gut microbes can be re-established in mice at the species level; among which, more than 1/3 have undergone significant changes (referred as to “variable microbes”), most of which were consistent across multiple human-mouse pairs and experimental settings. Consistently, one-third of human samples had changed their enterotypes, i.e. significant changes in their leading species after FMT. Mice fed with controlled diet showed significant decrease in the enterotype change rate (~25%) as compared those with non-controlled diet (~50%), suggesting a possible solution for rescue. Strikingly, most of the variable microbes have been implicated in human diseases, with some being recognized as causing species.

**Conclusions:** Our results highlighted the challenges of using mouse model in replicating human gut microbiota-associated phenotypes, provided useful information for researchers using mice in their gut microbiota studies and call for additional validations after FMT.

## Background

In recent years, it has been well established that alterations in gut microbiota can be linked to many aspects of human health and diseases [1–5], with many playing causal roles. For example, microbiota may play a fundamental role on the induction, training, and function of the host immune system, several studies have shown that gut microbes are associated with the occurrence and development of diseases [6–8]. However, establishing causal relationships between gut microbiota alterations and human phenotypes have proved to be difficult, despite advances in computational and experimental techniques [9].

Fecal microbiota transplant (FMT) of human fecal samples (fresh or frozen) to germfree (GF-) mice is one of the few available methods to establish causal relationships between human phenotypes and altered gut microbiota [10, 11]. Such “humanized” mice could be used to replicate human phenotypes at both physiological and molecular levels, study the relative contribution of the respective microbiota to host dysfunctions or disease phenotypes, and test the effects of the perturbation of certain species (often by addition of lab-cultured species into the mice) on the phenotypes of interests [12, 13]; they are thus also extremely valuable for finding possible intervention methods for human diseases. There have been numerous successful applications of such methods for the past a few years [14–17]. For example, recent studies have shown that the addition of *Bifidobacterium longum, Collinsella aerofaciens, and/or Enterococcus faecium* to GF-mice receiving FMTs from non-responding patients can greatly increase the efficacy of anti–PD-L1 therapy [18]. Similarly, age-associated differences in IgA responses can be repeated in young GF mice, those receiving FMTs from infant can demonstrate the influence of genetic factors on development of the gut microbiota and mucosal IgA responses [19].

However, engraftment of fecal microbial communities from human feces into germ-free mice results in only a partial resemblance to the donor microbiota [14], favoring those phylotypes adapted to the recipient species, due to the genetic, behavioral, physiological, and anatomical differences between the guts of mice and human [20, 21]. For example, mouse intestinal villi are taller than those of human, and intestinal pH is lower than human; mouse has a large cecum which is an important site for fermentation, while the human cecum is relatively small [20, 21]. Consequently, human and mouse native gut microbes are vastly different; for example, a recent study showed only 4% of human and mouse gut microbes were found to share 95% identity and a coverage of 90% [10, 11, 20, 22]. These results are consistent with the fact that most pathogens are able to infect multiple hosts but some are highly adapted to a single-host species [23].

Having becoming aware of the differences in human and mouse guts and their important implications, researchers have been taking measures to reduce their impacts on FMT, by either taking pre-experimental methods to select the most suitable model animals for studying certain bacteria, and/or feeding mice with human food [24]. The use of other models has also been considered recently [25]. Many studies have demonstrated that gut microbes can be (at least partially) influenced by diet [26]. For example, increased abundance of *Prevotella* is associated with increased dietary fiber in human [27]. These efforts are unfortunately not pain-free, and call for systematic analysis on the alterations in human fecal samples after transplanted to mice.

In this research we investigated the changes of human gut microbes after FMT by comparing abundances of the same species/genus in 1,713 human-mouse pairs. We focused on consistently changed species, i.e. those that are abundantly present in a significant proportion of human-mouse pairs with relative abundance higher than 0.1% and showed significant changes (with median |Log2FC| > 1; FC, fold change). We found strikingly over one-third human gut microbes have been significantly and consistently changed after FMT at both species and genus levels. Consequently, about one-third of human samples changed their enterotypes after FMT, i.e. significant changes in the leading species [28]. Strikingly, we found that most of the significantly changed species (referred to as “variable species” below) were implicated in human diseases (Tables 1 and S1). Human samples transplanted to recipient mice fed with human food (referred to as “controlled diet” group below) showed significant decreased in enterotype changes, suggesting possible methods of rescue; however, the latter may result in additional variable taxa. Our results would be informative to researchers that use (or plan to use) GF-mice in their gut microbiota studies. In addition, these results also call for additional validations of species of interests after FMT, e.g. to check if significantly changed species in different phenotype groups are still significantly changed after transplanted to FMT. So far, such validation has been mostly conveniently ignored.

**Table 1.**
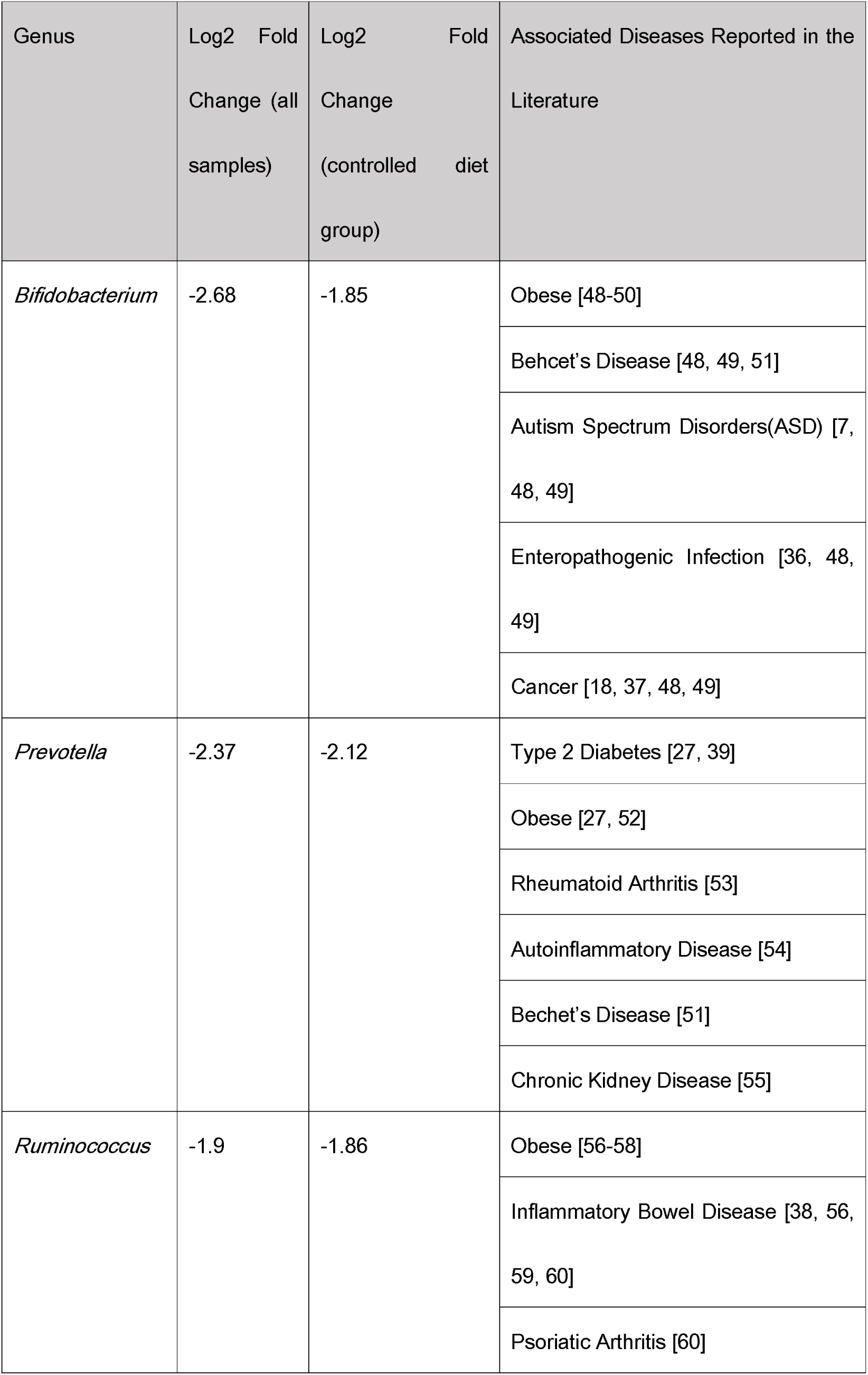
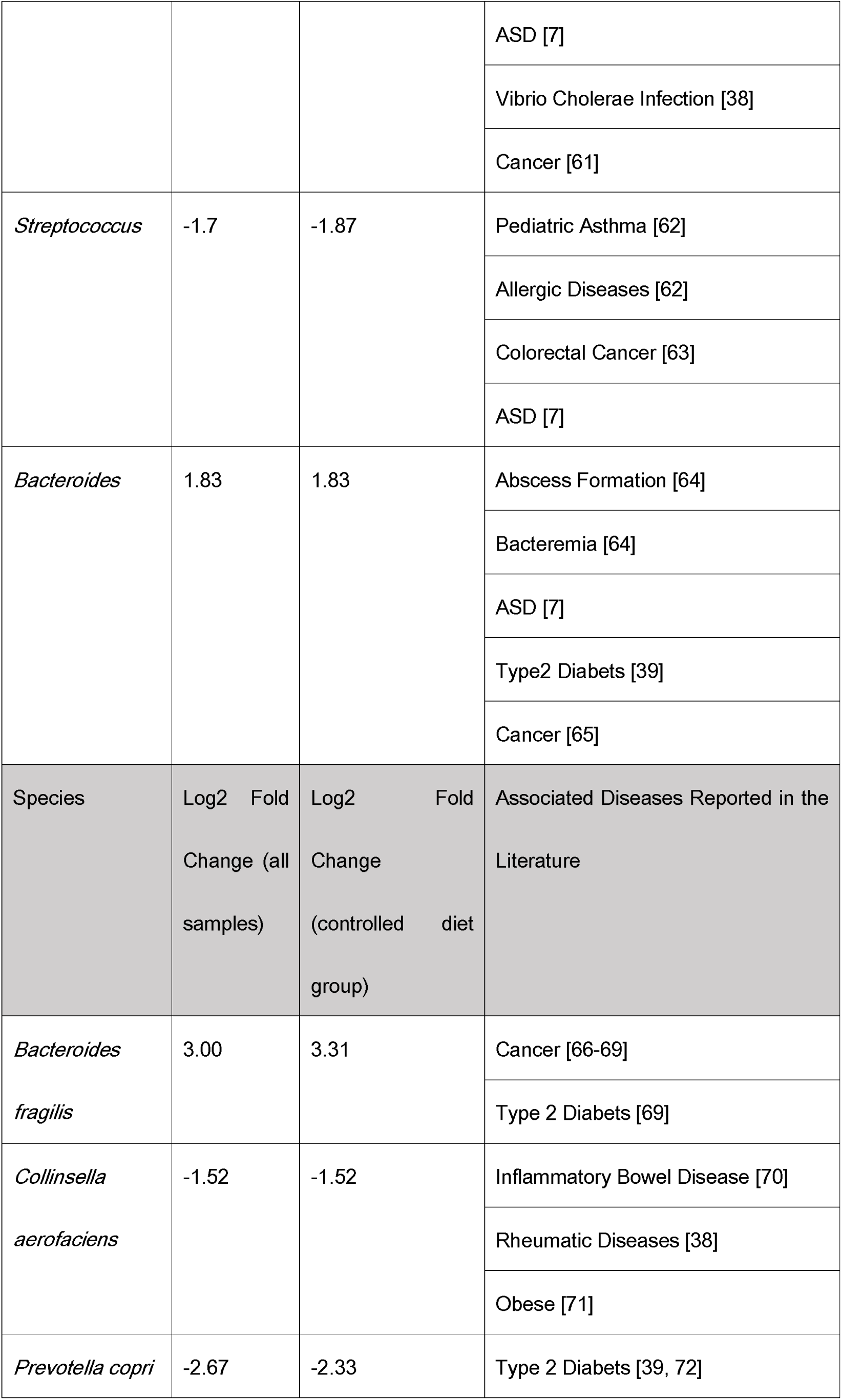
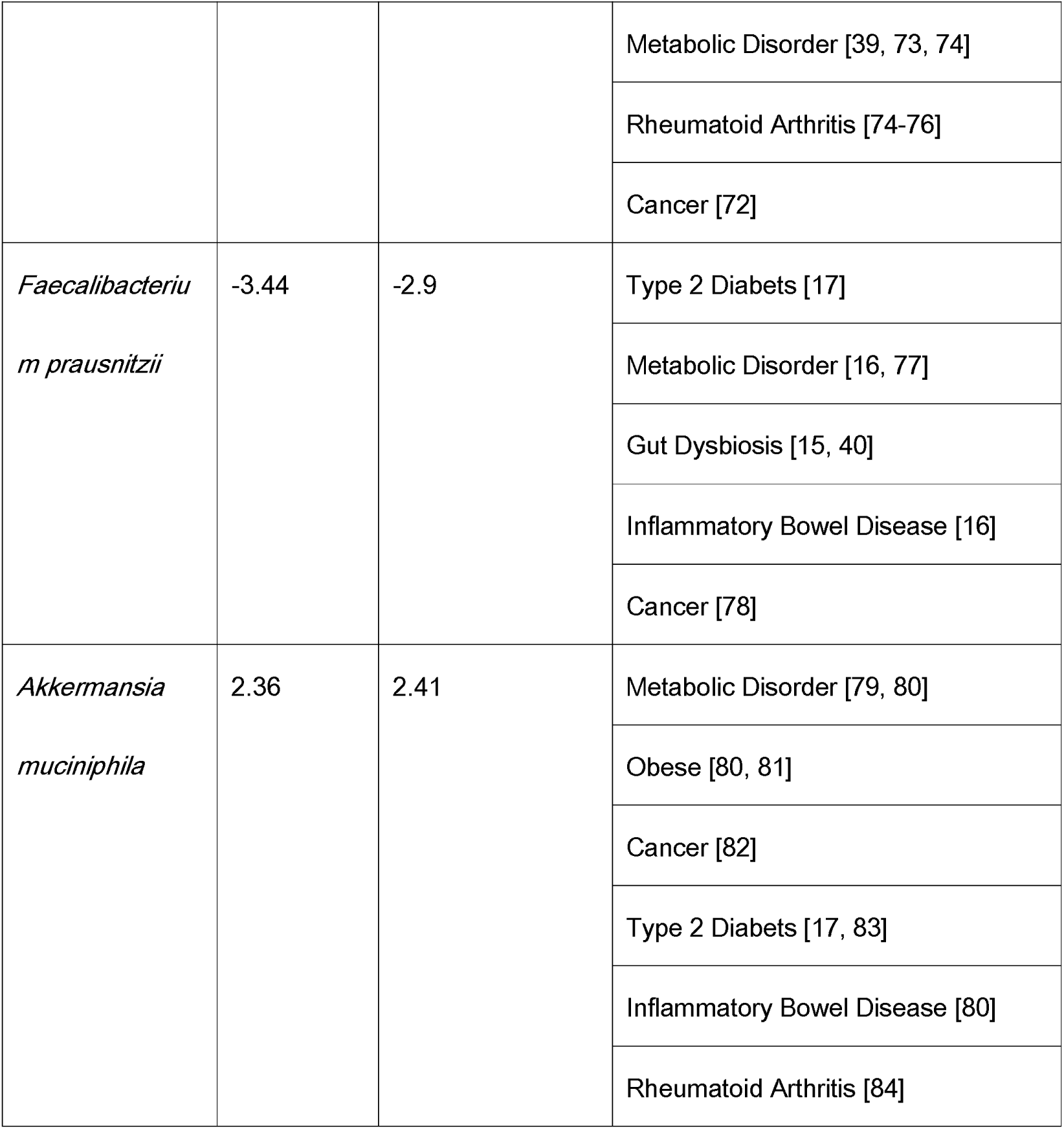
Selected variable taxa, their responses to diet changes and associations with human diseases.

## Results and discussion

### Less than 50% human gut microbes could re-establish in mice after FMT

We collected data from in total 1,713 human-to-mouse FMT experiments and analyzed the relative microbe abundances in fecal samples of the human donors and the corresponding germ-free mouse recipients (Materials and Methods). For each human-to-mouse pair, we computed the relative abundance changes before and after FMT. We first checked how many gut microbes could be re-established in mice after FMT. We limited our analysis to taxa that were supported by at least five sequencing reads in both samples of the human-mouse pairs. As shown in Figure 1, we estimated that on average only 47% human gut microbes could be re-established in mouse gut at species level after FMT; at genus level, ~60% could re-established. These numbers are consistent with some previous results [29] but differ significantly from others [30, 31]. For example, using 64 human-mouse pairs, Ridaura *et al* estimated 50~90% of human gut microbes at genus level could be re-established in mice after FMT, while others have found slightly higher re-establishment rates (>85% [30, 31]). However, they were all based on much smaller number of FMT experiments.

**Fig.1.**
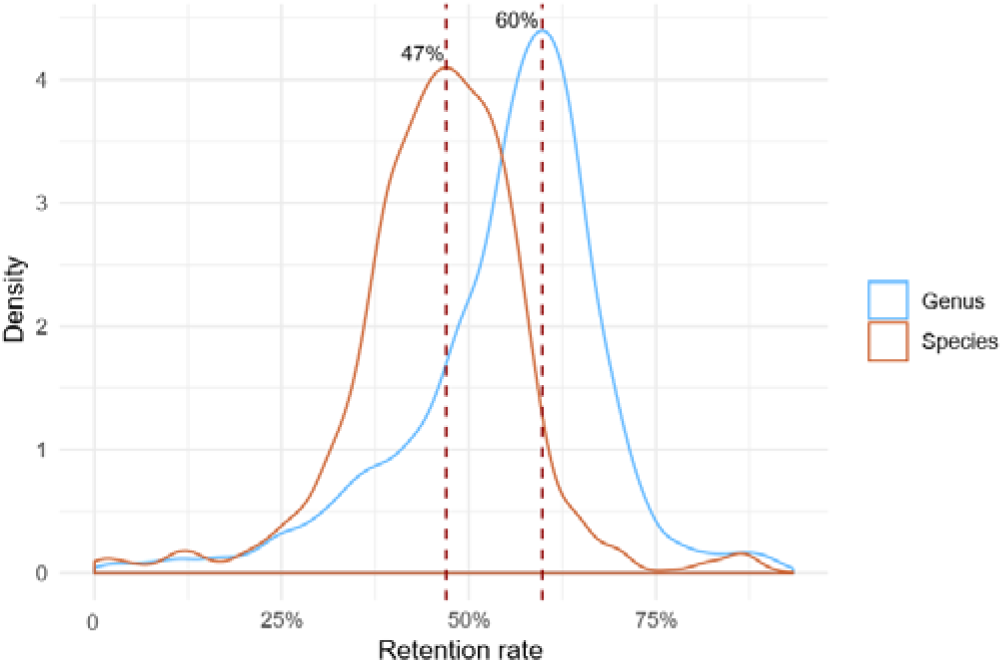
47% and 60% human gut microbes could be re-established in mouse gut after FMT at species (red) and genus (blue) levels respectively. We limited our analysis to taxa that were supported by at least five sequencing reads in both samples in each of the human-mouse pairs.

### Over one-third human gut microbes have been significantly and consistently changed after FMT to germ-free mice

To capture consistent changes in gut microbes we also required that a species or genus should be presented in at least five FMT pairs and two experimental conditions with minimal relative abundance of 0.1% in each sample (before and after FMT). We estimated that over one-third of the gut microbes have been significantly and consistently changed after the FMT. For example, at the genus level, we found that 37 out of in total 95 genera identified in our study been significantly changed after the FMT, i.e. their median abundance changes have been at least two-fold, accounting for 39.36% of the total genera (Figure 2). At species level, we found 84 (38.53%) out of 218 species have been significantly changed after FMT (Figure S1). We referred these significantly changed taxonomic groups as to “variable genera” and “variable species” respectively below, referred them together as to “variable taxa”; conversely, we referred others as to “stable taxa”. As shown in Figures 3 and S2, the variable taxa indeed showed consistent changes across experimental conditions.

**Fig.2.**
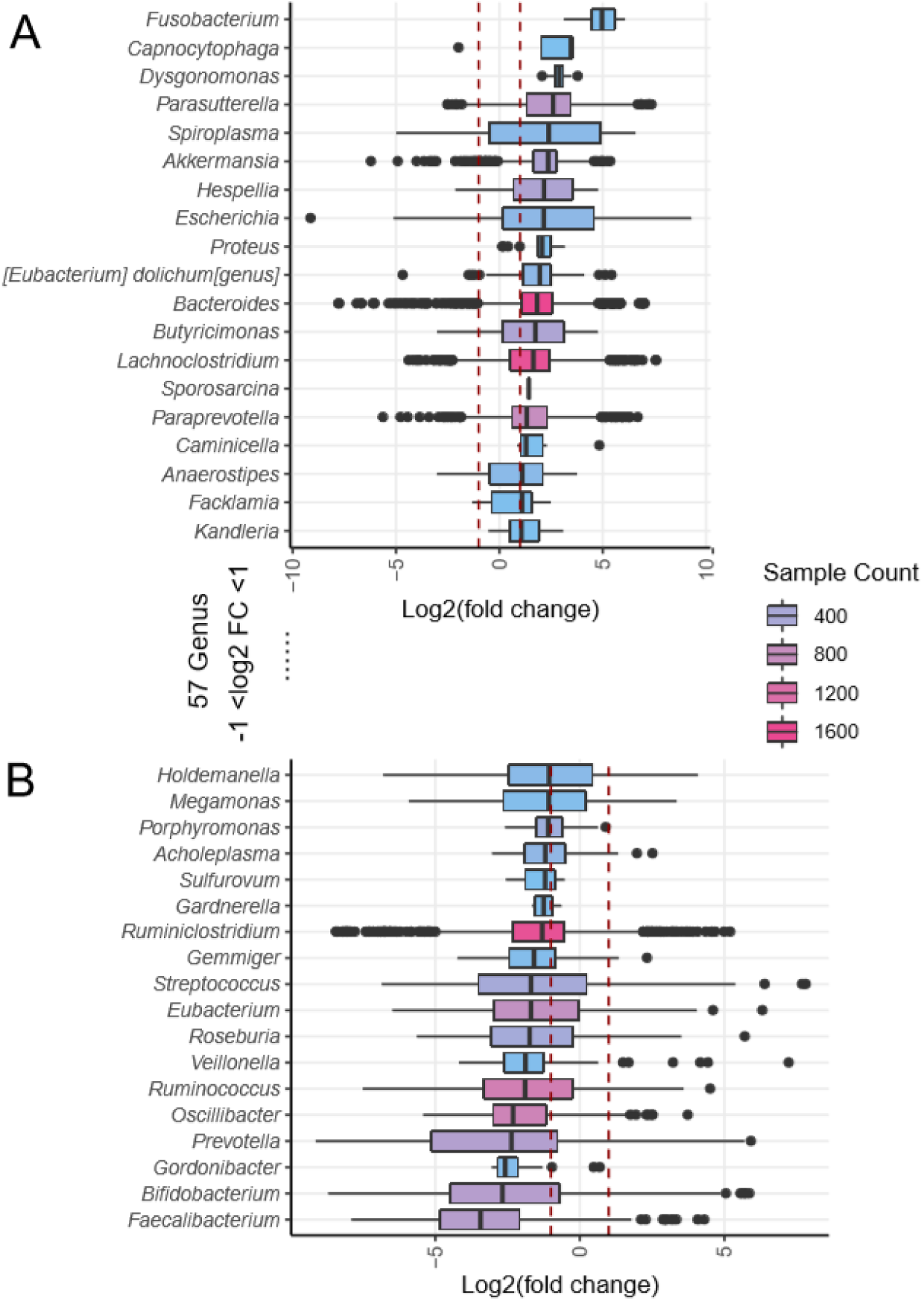
Over one-third of the genera of human gut microbes were significantly and consistently altered after transplanted to germ-free mice. The box plot shows the relative abundances changes (log2-transformed; Log2FC) at genus level in the 1,713 human-mouse pairs we have collected (see Materials and Methods for details). Boxes were colored according to the number of samples (sample count) in which a genus was found; sample counts in this plot range from 81 to 1,713. If the median Log2FC value of a genus is higher (panel A) or lower (panel B) than 1, it is considered to be significantly altered after FMT.

**Supplementary Fig.1.**
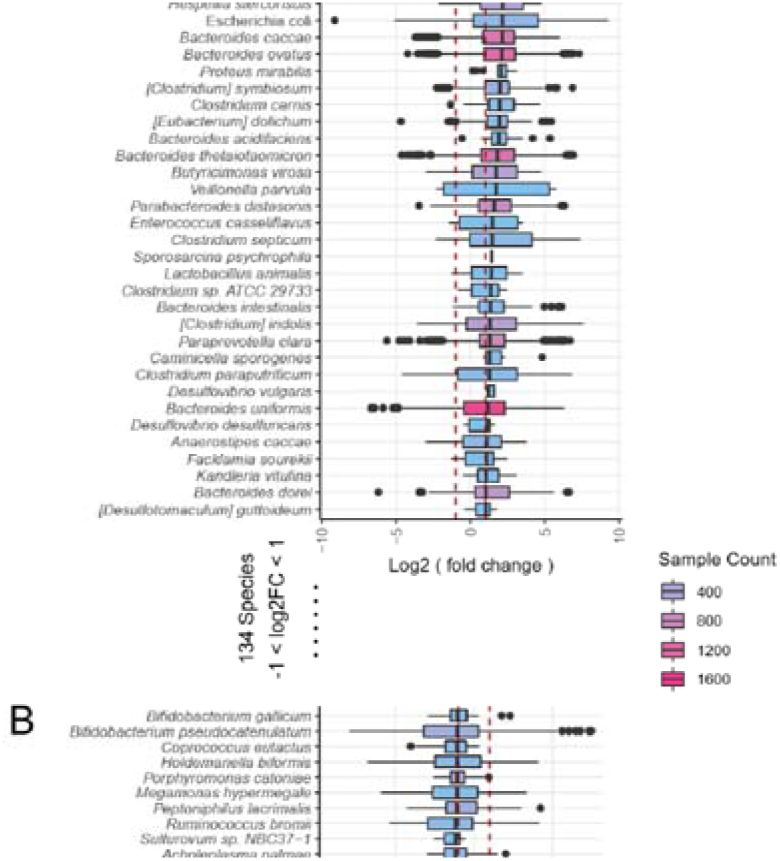
Over one-third of the species of human gut microbes were significantly and consistently altered after transplanted to mice. The box plot shows the relative abundances changes (log2-transformed; Log2FC) at species level in the 1,713 human-mouse pairs of samples we have collected. Boxes were colored according to the number of samples (sample count) in which a genus was found; sample counts in this plot range from 26 to 1,697. If the median Log2FC value of a species is higher (panel A) or lower (panel B) than 1, it is considered to be significantly altered after FMT.

**Fig.3.**
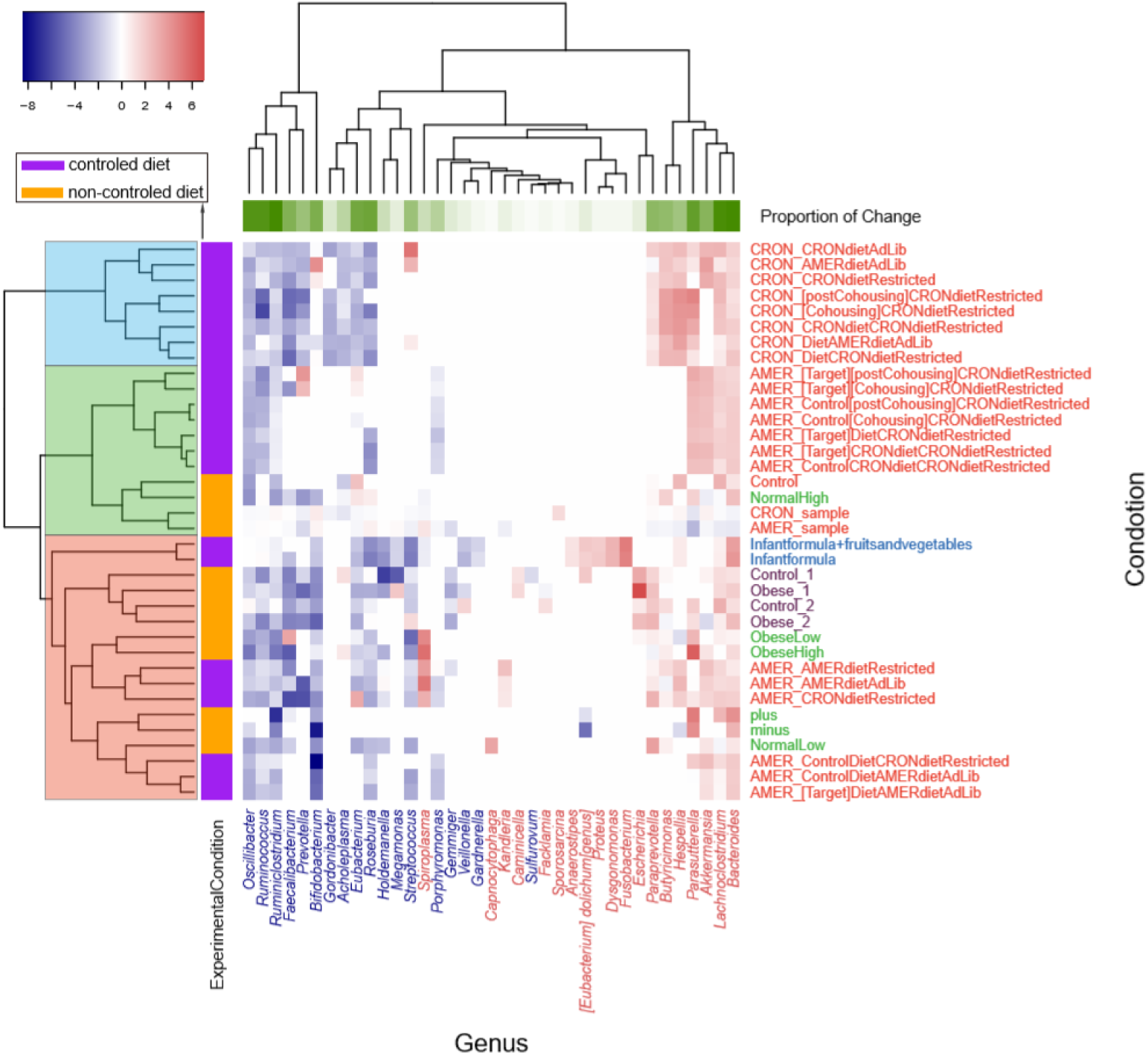
Variable genera showed consistent changes across experiment conditions. Shown here is the median fold changes (log2 transformed; Log2FC) at genus level in samples of the same experiment conditions, i.e. for each of the 33 experiment conditions, we calculated the median Log2FC of each genus in human-mouse pairs, and plotted the results as a heatmap. The experimental conditions from the same studies are marked with the same color (Y-axis labels on the right). Significantly increased and decreased genera are marked in red and blue respectively (X-axis labels). The heatmap is colored according to the Log2FC values. Rowside colors are used to show whether controlled diets were used in the experiments, while column-side colors are used to show proportion of experimental conditions in which the genus has been significantly changed.

**Supplementary Fig.2.**
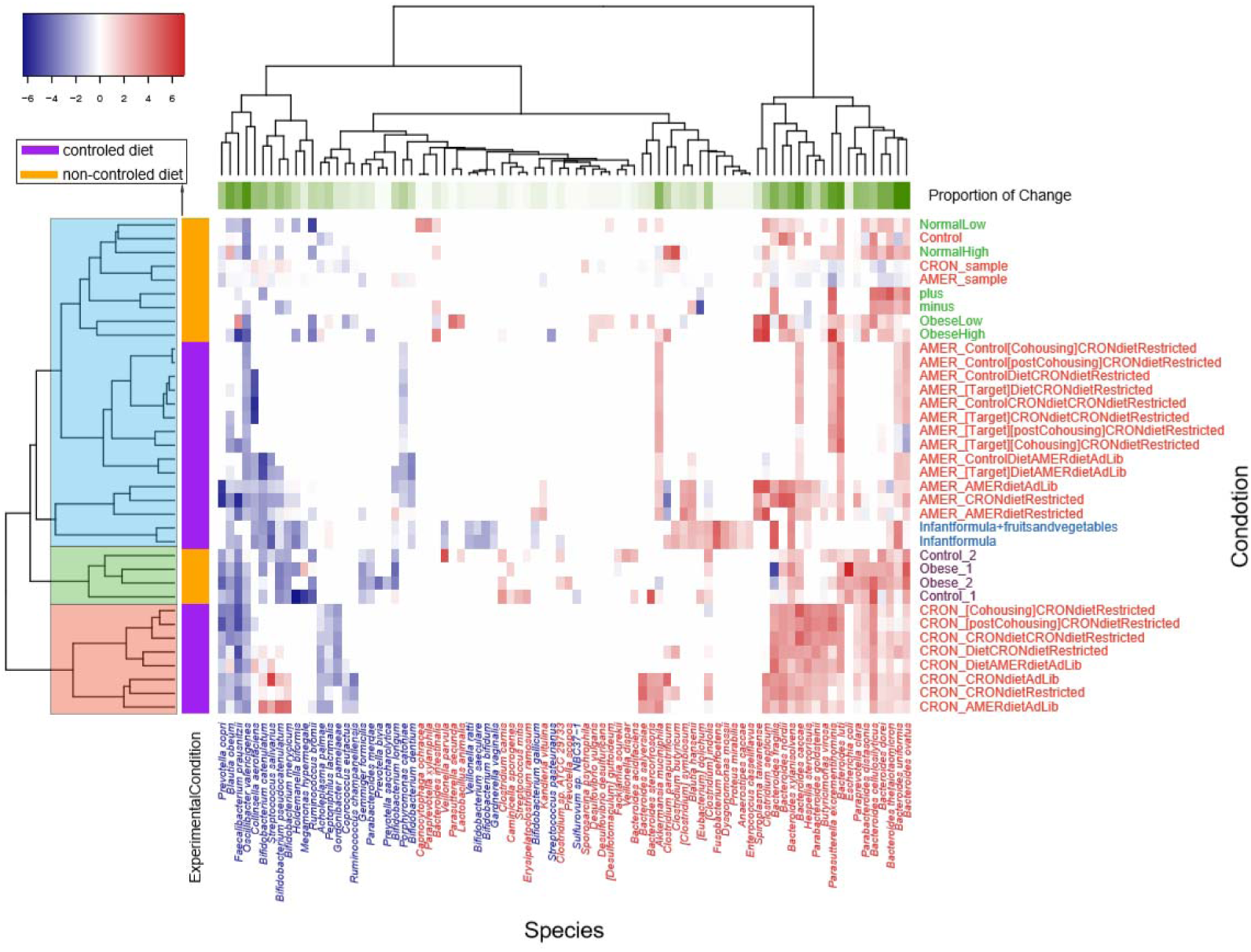
Variable species showed consistent changes across experiment conditions. This figure is similar to Figure 2, only that the results of variable species are plotted. Shown here is the median fold changes (log2 transformed; Log2FC) at species level in samples of the same experiment conditions, i.e. for each of the 33 experiment conditions, we calculated the median Log2FC of each species in human-mouse pairs, and plotted the results as a heatmap. The experimental conditions from the same studies are marked with the same color (Y-axis labels on the right). Significantly increased and decreased species are marked in red and blue respectively (X-axis labels). The heatmap is colored according to the Log2FC. Rowside colors are used to show whether controlled diets were used in the experiments, while column-side colors are used to show proportion of experimental conditions in which the genus has been significantly changed.

### Changes in enterotypes were relatively stable over time but significantly affected by diet

We also checked the changes in enterotypes after FMT. Human gut microbes could be classified into three enterotypes, each with distinct leading species [28]. Our results showed that the mixed data from humans and mice can also form three enterotypes, as shown in Figure S3. Although the concept of enterotypes has been hotly debated recently, they are nevertheless is a useful approach for a better understanding of complex biological problems [32]. Changes from one enterotype to the other often indicate significant alterations in overall gut microbe profiles [28]. We calculated the enterotypes for all fecal samples before and after FMT using a method previously described (http://enterotype.embl.de/enterotypes.html); the overall classification of all samples is shown in Figure S3. Strikingly, we found that 32.17% of the human samples changed their enterotypes after FMT (Figure 2A). These results are consistent with the results that 1/3 of the species and/or genera have been significantly altered after FMT. For example, the top three driving genera of enterotype I, namely *Bifidobacterium, Clostridiodes* and *Dysgonomonas* increased significantly after FMT (Figure 2). For enterotype II, two of the top three driving genera, namely *Alistipes* and *Bacteroides* increased significantly after FMT, while *Lactonifactor*. Similarly, the top three driving genera of enterotype III, namely *Butyricimonas, Odoribacter* and *Paraprevotella* were all significantly changed, although with different trends. As shown in Figure S4, 57.2% of type I samples changed their enterotypes after FMT, followed by type II (31.2%%) and then type III (24.3%).

**Supplementary Fig.3.**
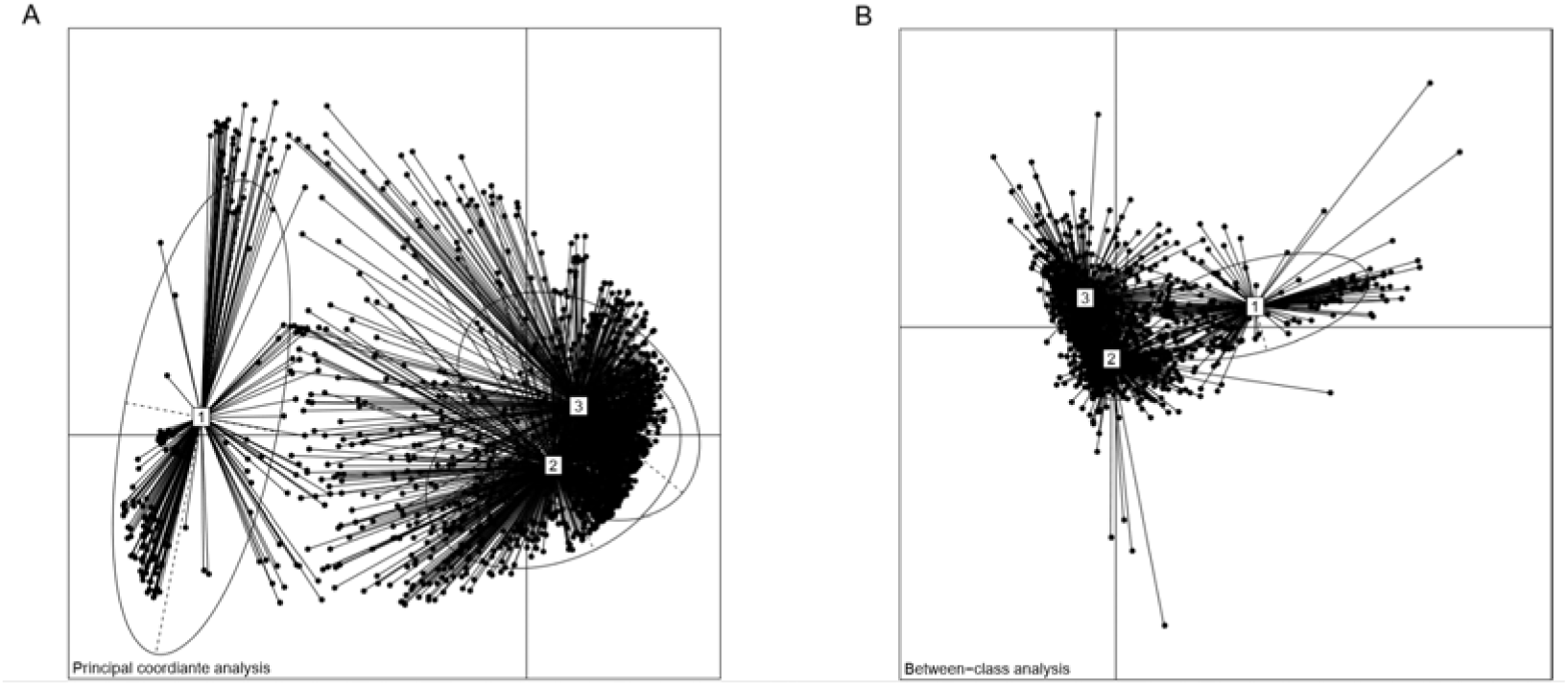
Enterotypes of all human fecal samples sequenced before and after FMTs. **A.** Principal coordinates analysis (PCoA), **B.** Between-class analysis (BCA).

**Supplementary Fig.4.**
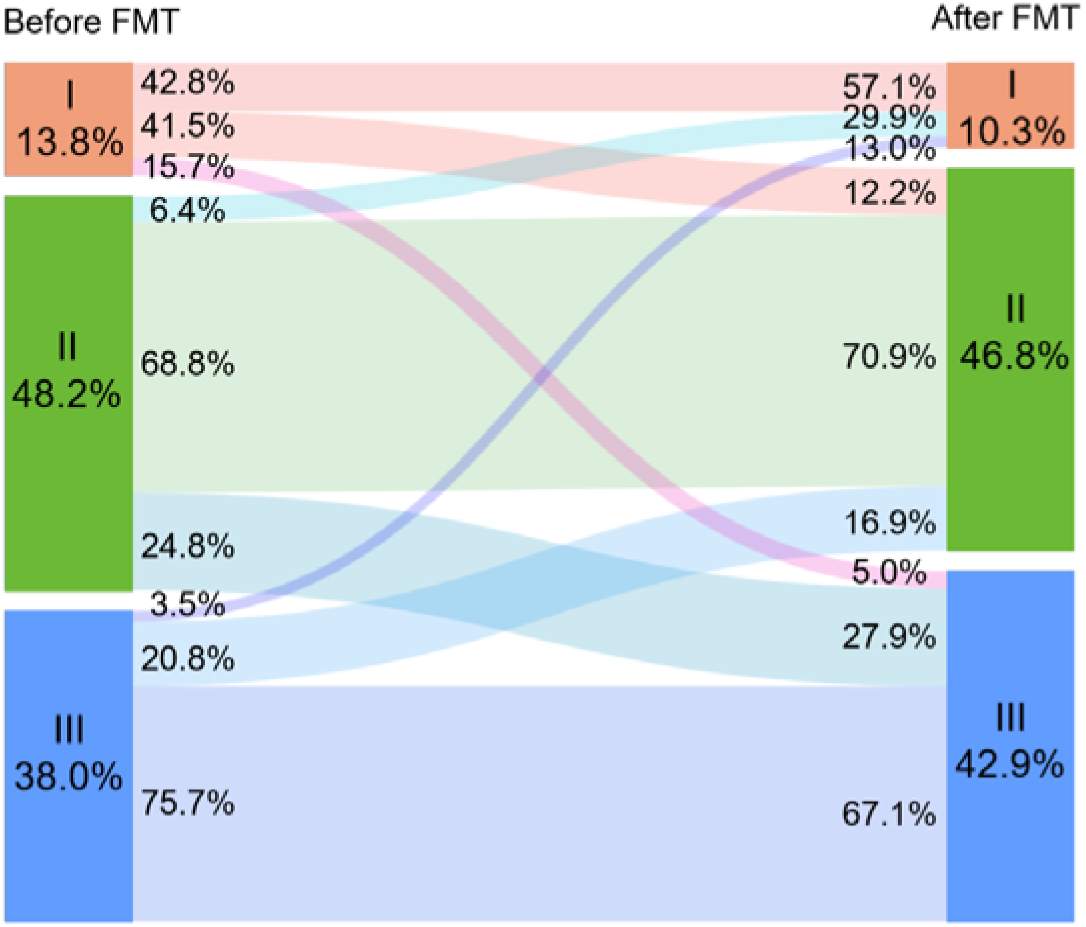
Enterotype changes in samples whose original enterotypes are I, II and III before FMT. Enterotype changes in all samples, i.e. recipient mice were fed with both human food and mouse food.

We next examined factors contributing to the enterotype changes. Previous studies have revealed that diet has a great influence on intestinal microbes [33]. After divided the human-mouse pairs into two groups according to whether the mice were fed with a controlled diet (i.e. human food, see Materials and Methods for details), we found that mice with controlled diet showed significant decrease in the enterotype change rate (23.52%) as compared those with non-controlled diet (48.97%). As shown in Figure 4B and 4C, in the non-controlled diet group, ~95.6%, 49.2% and 26.8% of the human samples of enterotypes I, II and III changed their enterotypes after transplanted to mice, respectively. These numbers decreased to 33.1%, 21.2% and 23.2% respectively in the controlled diet group, largely due to decreased changes in the leading taxa of the respective enterotypes (Table S1). These results are consistent with previous results that diets have significant impact on gut microbiota [33, 34], and may provide useful hints on the better retention of human gut microbiota in the humanized mice. However, we found that controlled diet may induce additional variable taxa at both species and genus levels (Figure S5), indicating complex reactions of mouse gut microbiota to human diets, and limited capacity of controlled diets on recipient mice.

**Fig.4.**
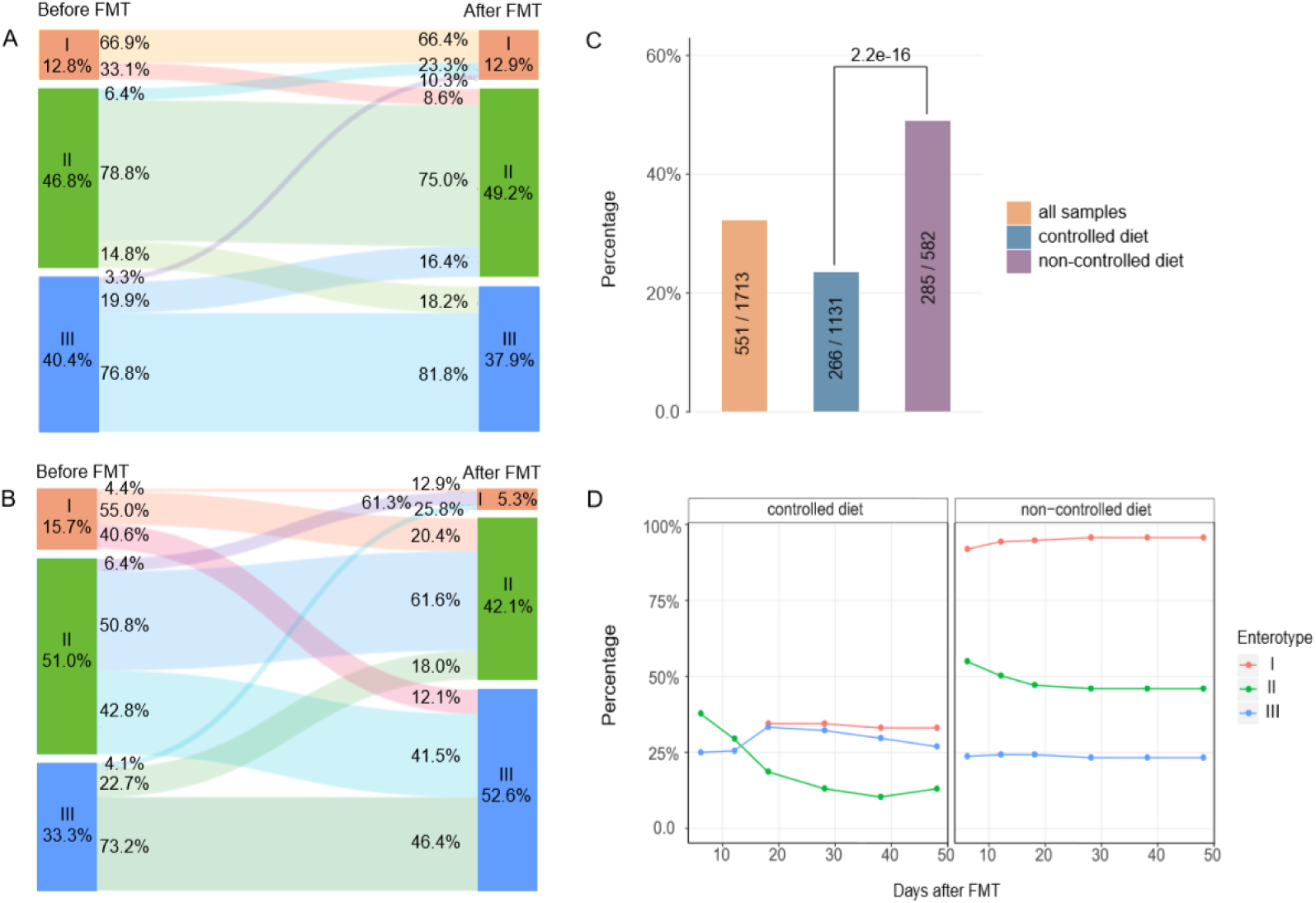
A significant proportion of samples changed their enterotypes after FMT, such change is significantly affected by diets of recipient mice. **A**, enterotype changes in the “controlled diet” group, i.e. recipient mice were fed with human food. **B**, enterotype changes in the “uncontrolled diet” group, i.e. recipient mice were fed with mouse food. **C**, Overall enterotype change rates of all samples (left) and samples in the “controlled diet” group (middle) and “non-controlled diet” group (right); enterotype changes have been significantly decreased in the “controlled diet” group. **D**, Enterotype changes in groups with the two diet types as a function of time (days after FMT) at which fecal samples were sequenced.

**Supplementary Fig.5.**
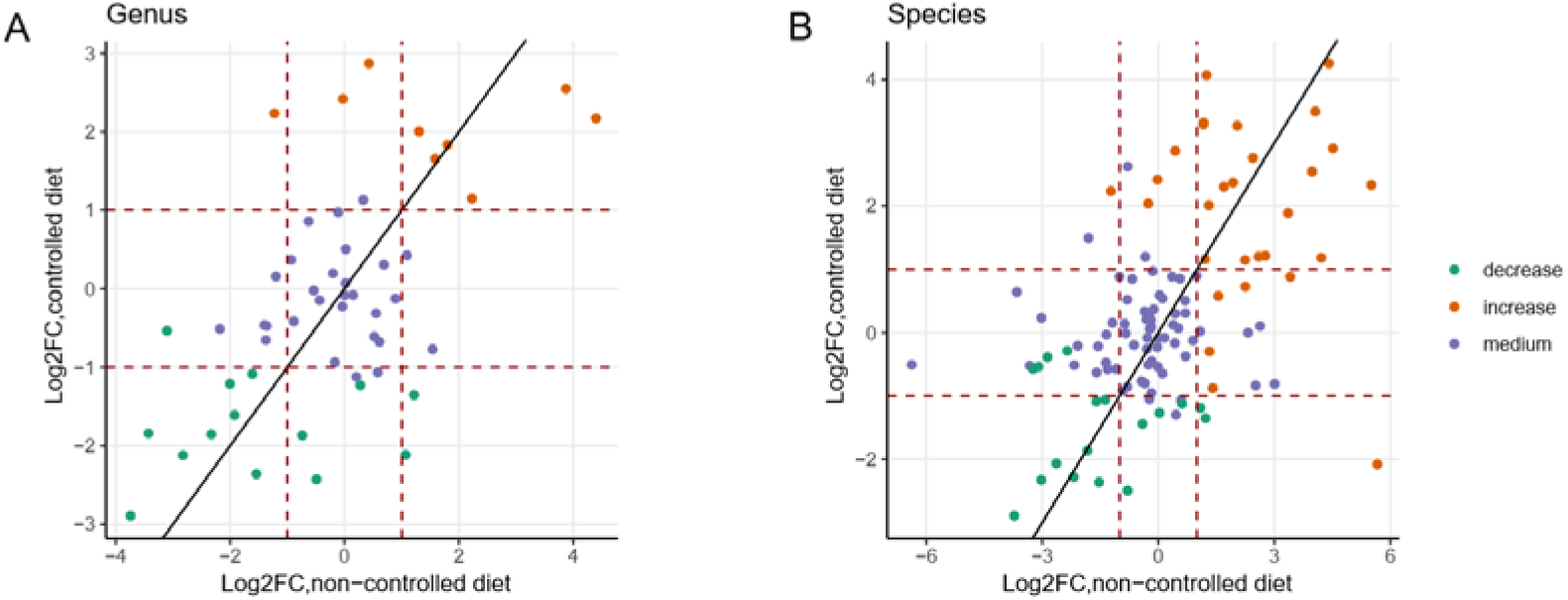
Complex interactions of mouse gut microbiota to human diets. As compared to non-controlled diet, controlled diet could reduce differences in human microbiota at both genus (A) and species (B) levels after FMT; however, the latter may result in additional variable taxa. Each dot represents a genus/species; green and red indicate variable taxa in Figure 2, while blue indicate non-variable taxa.

We also checked whether changes in the gut profiles could be different at different time points after translated to mice. Surprisingly, dividing human-mouse pairs into subgroups according to the days after FMTs, we found that the enterotype change rate was the relatively stable over time, with the exception of human samples with initial enterotype II (the green lines in Figure 4D), which showed the highest rates of enterotype change at the first 10~20 days; we found similar trends in mice with controlled and non-controlled diets (Figure 4D). These results suggest a possible “shock” period immediate after by transplants of the fecal microbiota, followed by adaptation and a stable state [19, 35].

### Distinct changes in species of the same genus

We next sought to check whether or not species in the same genus tend to have similar trends after FMTs. As shown in Figure 5, we identified four typical patterns among in total 27 genera that contained multiple species. Pattern one (Figure 5A and S6A) consists of nine genera; in this group, all species and very often the corresponding genus are “stable taxa”. Pattern two (Figure 5B and S6B) also consists of five stable genera. However, they included both increased- and decreased-variable species, sometimes also stable species; the overall abundances of the genera were not changed after FMT. Pattern three (Figure 5C and S6C) and four (Figure 5D and S6D) consists of five increased- and eight decreased-“variable genera” respectively; their included species are either increased/decreased variable-species, or stable species. They account for 33.33%, 18.52%, 18.52% and 29.63% of the multi-species genera respectively. These results indicated distinct species preferences within the genera between human and mouse.

**Fig.5.**
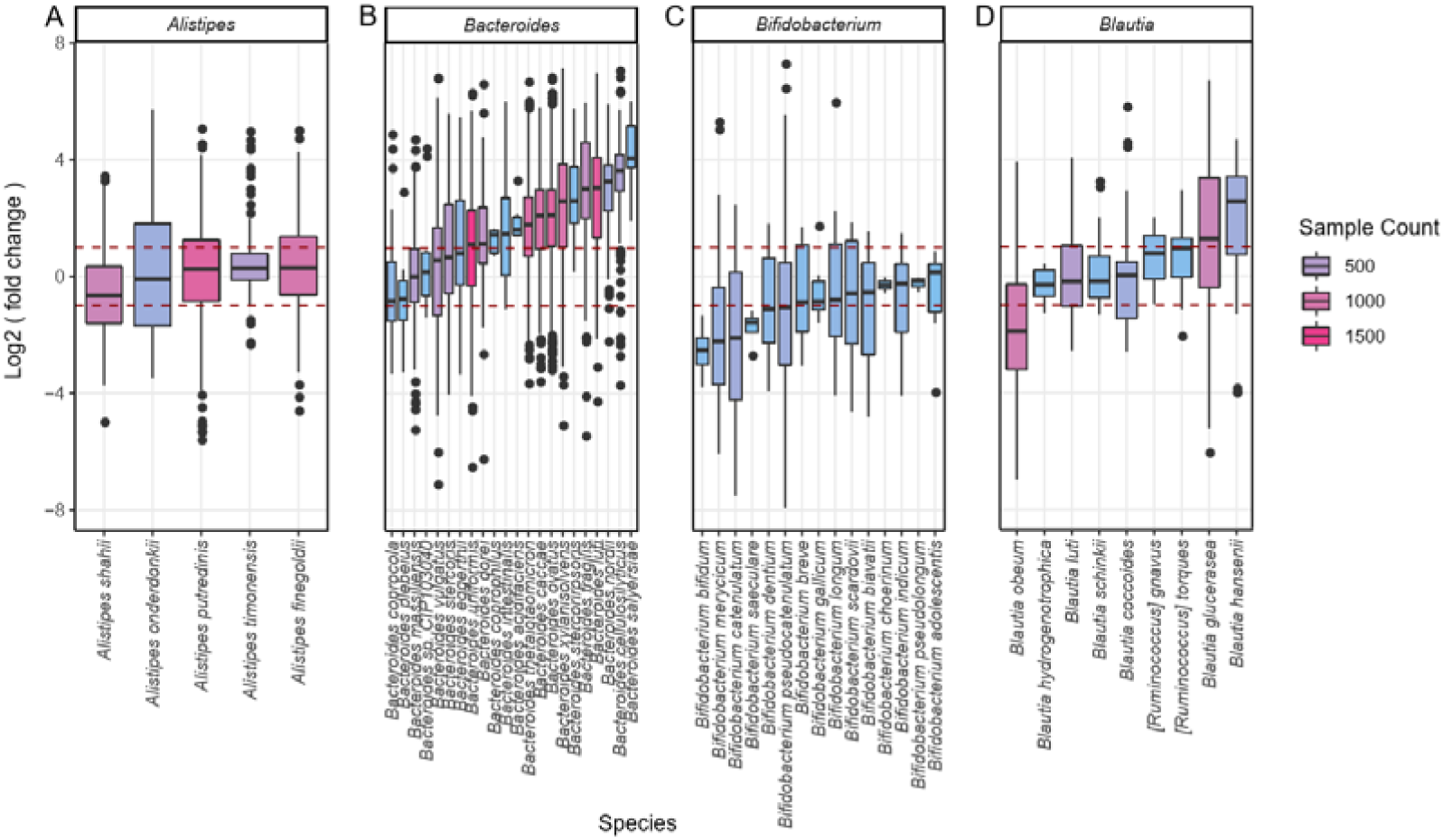
Species of the same genus may undergo distinct changes after FMT. Four typical genus types of change. The vertical axis is the ratio of human donor abundance to post-transplant rate of change, and the horizontal axis is the different species under the same genus.

**Supplementary Fig.6.**
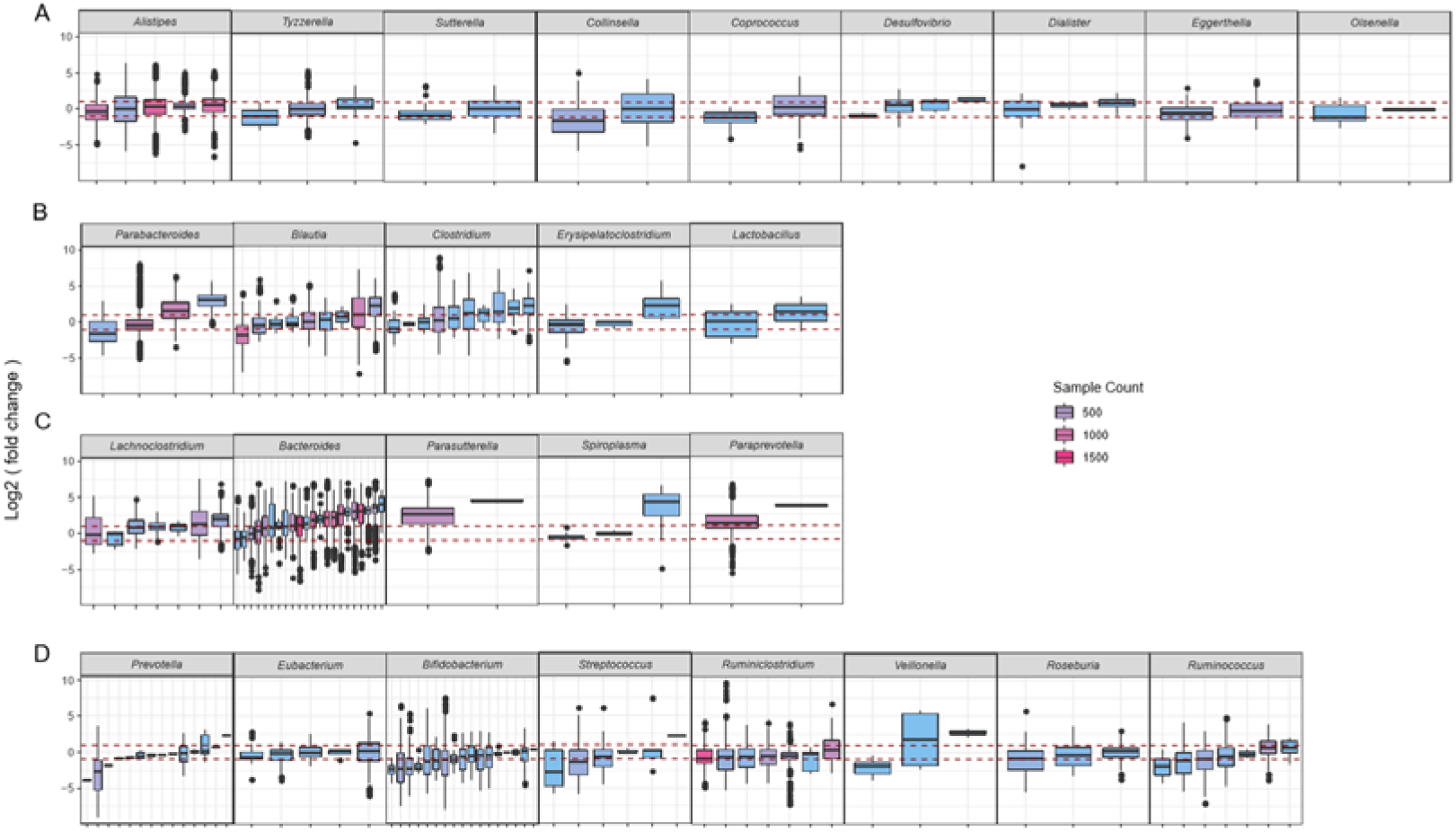
Four types of multi-species genus with distinct change patterns at species levels. **A** stable genus whose member species are all stable species. **B** stable genera whose member species also include variable species. **C** variable genera significantly increased in abundances after FMT whose member species are either stable- or increased variable-species. **D** variable genera significantly decreased in abundances after FMT whose member species are either stable- or decreased variable-species.

### Most variable taxa are linked to human health and diseases

Strikingly, we found most of the identified variable taxa were previously implicated in various human diseases and/or can be used in disease intervention and treatment (Table 1 and S1). Shown in Figure 6 are eight selected variable taxa (four genera and four species) that were relatively well studied in human diseases. Among which, *Bifidobacterium* has been implicated in obese, and can protect humans from enteropathogenic infection through production of acetate [36]; recently, *Bifidobacterium*, *Akkermansia muciniphila* and other species were found to be able promote antitumor immunity and facilitates anti-PD-L1 efficacy [37]. *Ruminococcus obeum* was shown to be able to competitively inhibit and control the infection of *Vibrio cholerae* [38]. *Prevotella copri* can induce insulin resistance, aggravate glucose intolerance and augment circulating levels of branched-chain amino acids [39]. *Faecalibacterium prausnitzii* exhibits anti-inflammatory effects on cellular and [2,4,6trinitrobenzenesulphonic acid (TNBS)-induced] colitis models, partly due to secreted metabolites able to block NF-ΚB activation and IL-8 production [40].

**Fig.6.**
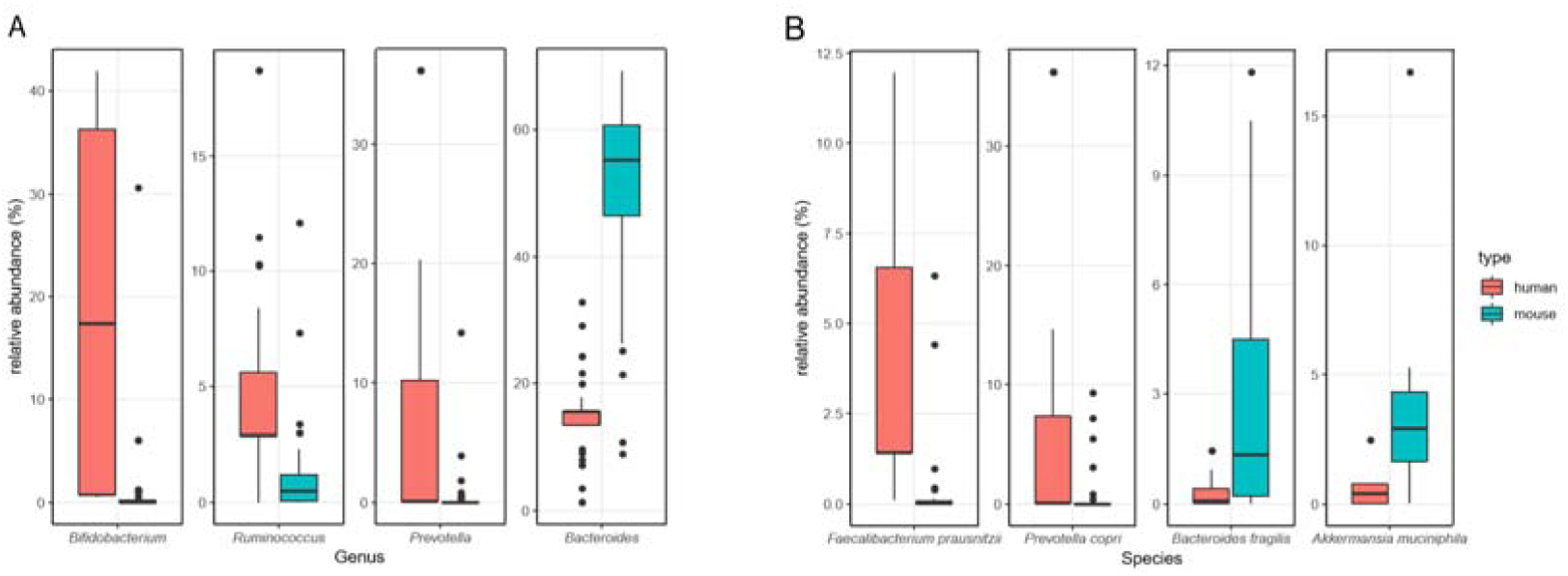
Exemplary variable genus (A) and species (B) that showed significant changes before and after FMTs. Boxplots show their relative abundances in human (before FMT, red) and mouse (after FMT, blue). Please consult Table 1 and S1 for more details of their associations with human diseases; please consult Figures S7 and S8 for all the variable genera and species that showed significant changes before and after FMTs.

**Supplementary Fig.7.**
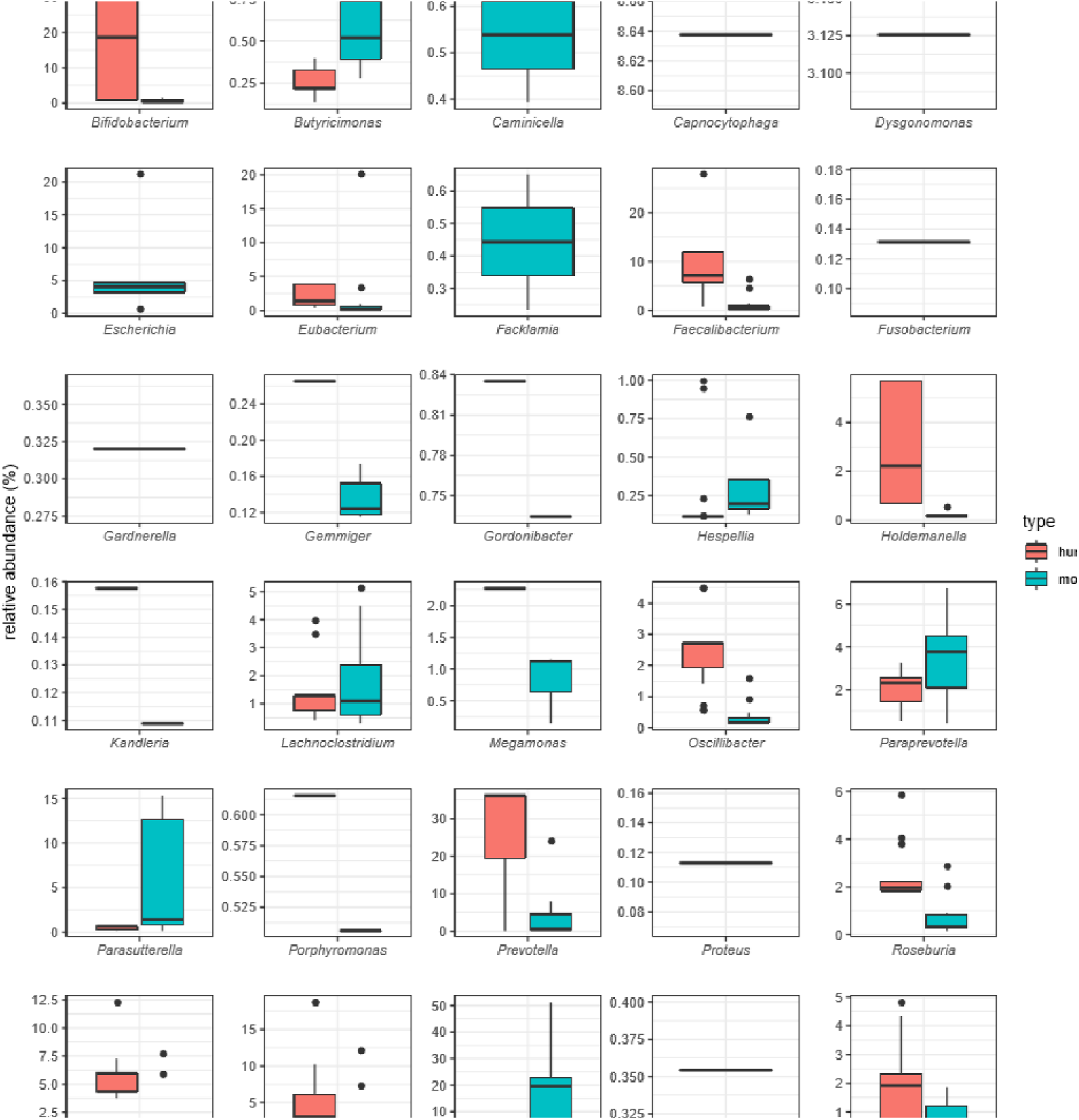
Variable genera and their relative abundances in human (before FMT) and mouse (after FMT). All genera with |Log2FC| > 1 were shown here. Please consult Table 1 and S1 for more details of their associations with human diseases.

**Supplementary Fig.8.**
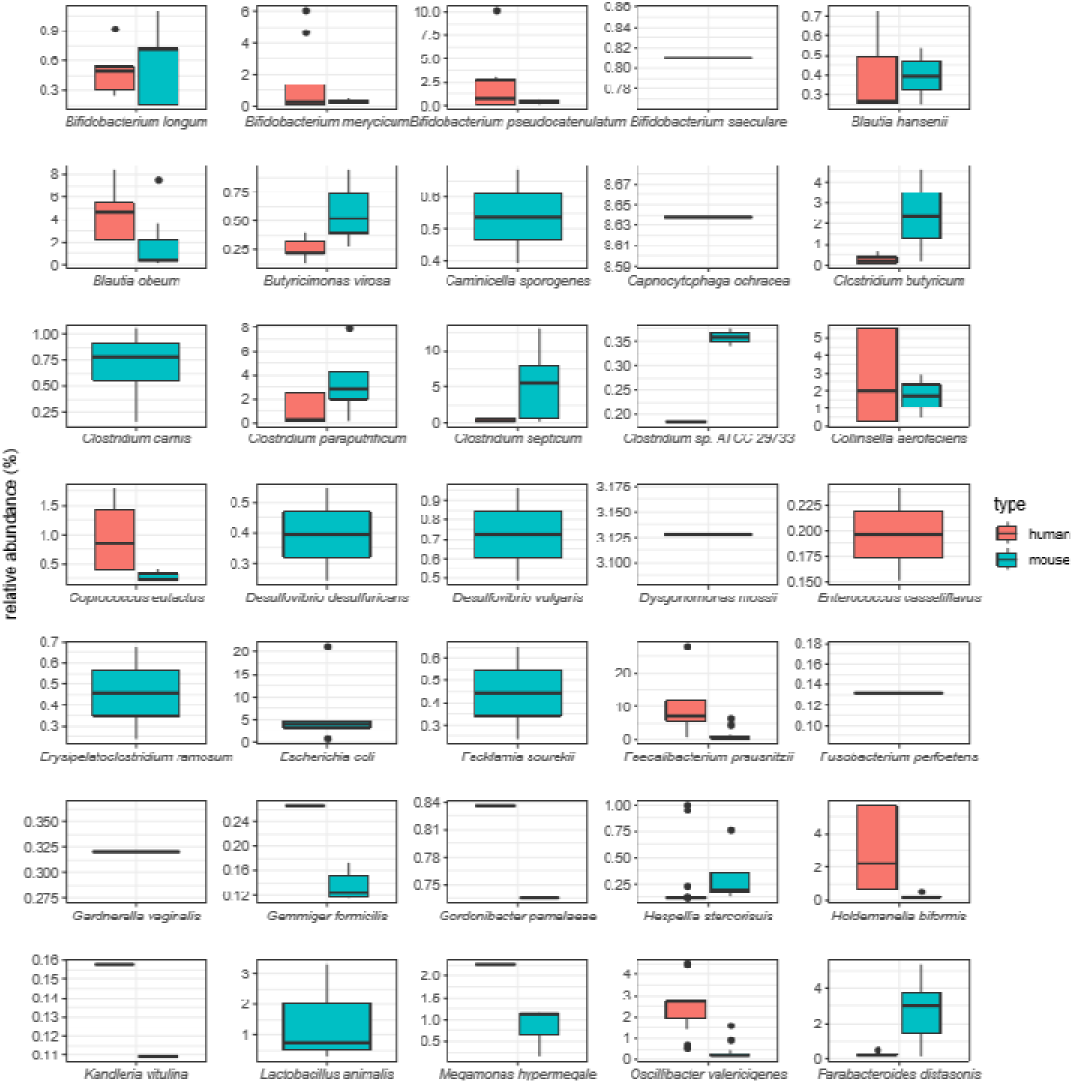
Variable species and their relative abundances in human (before FMT) and mouse (after FMT). All species with |Log2FC| > 1 were shown here. Please consult Table 1 and S1 for their association with human diseases.

These results highlight the challenges in the use of the mouse model in gut microbiota studies: not only it will be difficult have similar human phenotypes (at both philological and molecular levels) in recipient mice if the suspected causal species would be significantly decreased in abundances after FMT, but also that it would difficult to transfer the treatment methods back to human if the interventional species adapts much better to the mouse but than that of the humans (e.g. *Akkermansia muciniphila*).

## Conclusions

Mice have been widely used in human disease studies and are one of the few available experimental methods for establishing causal relationships between altered microbial abundances in human gut and diseases, and finding intervention and treatment methods for gut-microbiota related diseases. However, due to intrinsic differences between mouse and human guts, transplantation of fecal microbial communities from human feces into GF-mice could only re-establish part of the donor microbiota, most of which are those had been adapted to the recipient species [14]. Despite widespread awareness of the differences and a handful of differential species identified in small-scale studies, a systematic analysis on the alterations in human fecal samples after transplanted to mice is yet to be conducted.

In this study we collected and analyzed more than 1,713 FMT experiments, and identified microbial abundance changes in before and after the transplantation (hence referred to as human-mouse pairs). We focused on species that are abundantly present in more than 400 human-mouse pairs with relative abundance higher than 0.1% and showed significant changes (with median |Log2FC| > 1; FC, fold change); these changes are thus more likely to be consistent across experimental conditions. Strikingly, we found over one-third human gut microbes have been significantly and consistently changed after FMT at both species and genus levels, including those that are leading species in enterotype analysis. Human feces transplanted to recipient mice fed with human food (the “controlled diet” group) showed significant decreased in enterotype changes, suggesting a possible method for reduce such differences; however, controlled diet may induce additional variable taxa at both species and genus levels (Figure S5), indicating complex reactions of mouse gut microbiota to human diets, and limited capacity of controlled diets on recipient mice.

These results highlighted the challenges of selecting mice as the animal models in gut microbiota studies. That is, it will be difficult to replicate human phenotypes that we thought to be caused by certain increased microbes, if the suspected species are to be significantly decreased in abundances after FMT solely due to the differences in human and mouse intestines. In addition, when transferring findings in mice back to humans, it would difficult if the interventional species adapts much better to the mouse but than that of the humans.

More strikingly, most of the variable taxa were implicated in human diseases. Our results thus would be informative to researchers that use (or plan to use) GF-mice in their gut microbiota and disease association studies. In addition, our results also call for additional validations of species of interests after FMT. For example, researchers should also check if significantly changed species in different human phenotype groups are still significantly changed after transplanted to FMT. In other words, fecal samples from both human patients and healthy controls should be transplanted to GF-mice; researchers should then be concerned with not only if the phenotypes of interests are replicated in mice, but also if the differentially abundant taxa in the GF-mice receiving patient and healthy feces are the same ones found in human samples. So far, such validation has been mostly conveniently ignored.

## Methods

### Data

We performed an extensive search in PubMed and NCBI SRA database to search for publications and/or deposited metagenomic sequencing data from human-to-mouse FMT experiments. Since the number of published data of the relevant experiments is relatively small, we used two keywords “microbiota transplanted into mice” and “human fecal microbiome mice” to expand the range of searching results. We limited our search for studies published in the last five years (from Jan 01, 2014 to date). We obtained in total 58 studies, however among which only 13 were human to mouse FMTs (Figure S9). We excluded experiments in which the recipient mice were genetically modified, not germ-free, or treated with antibiotics; we also excluded experiments in which the human donors were treated with antibiotics, or only a few (less than ten) human donors were used (Figure S9). In the end, we found four studies that met our searching criteria (Table 2), and all of which used 16S sequencing to survey gut microbiota before and after FMTs. These studies included in total 1,713 human-mouse pairs. We downloaded 16S DNA sequencing data from the NCBI SRA database using a command-line tool fastq-dump of the SRA-tools (https://github.com/ncbi/sra-tools, accessed in July 2018). We obtained the related meta-data, including human-mouse pairing, experimental conditions, dates of sampling after FMT from the corresponding publications and/or the NCBI SRA database.

**Supplementary Fig.9.**
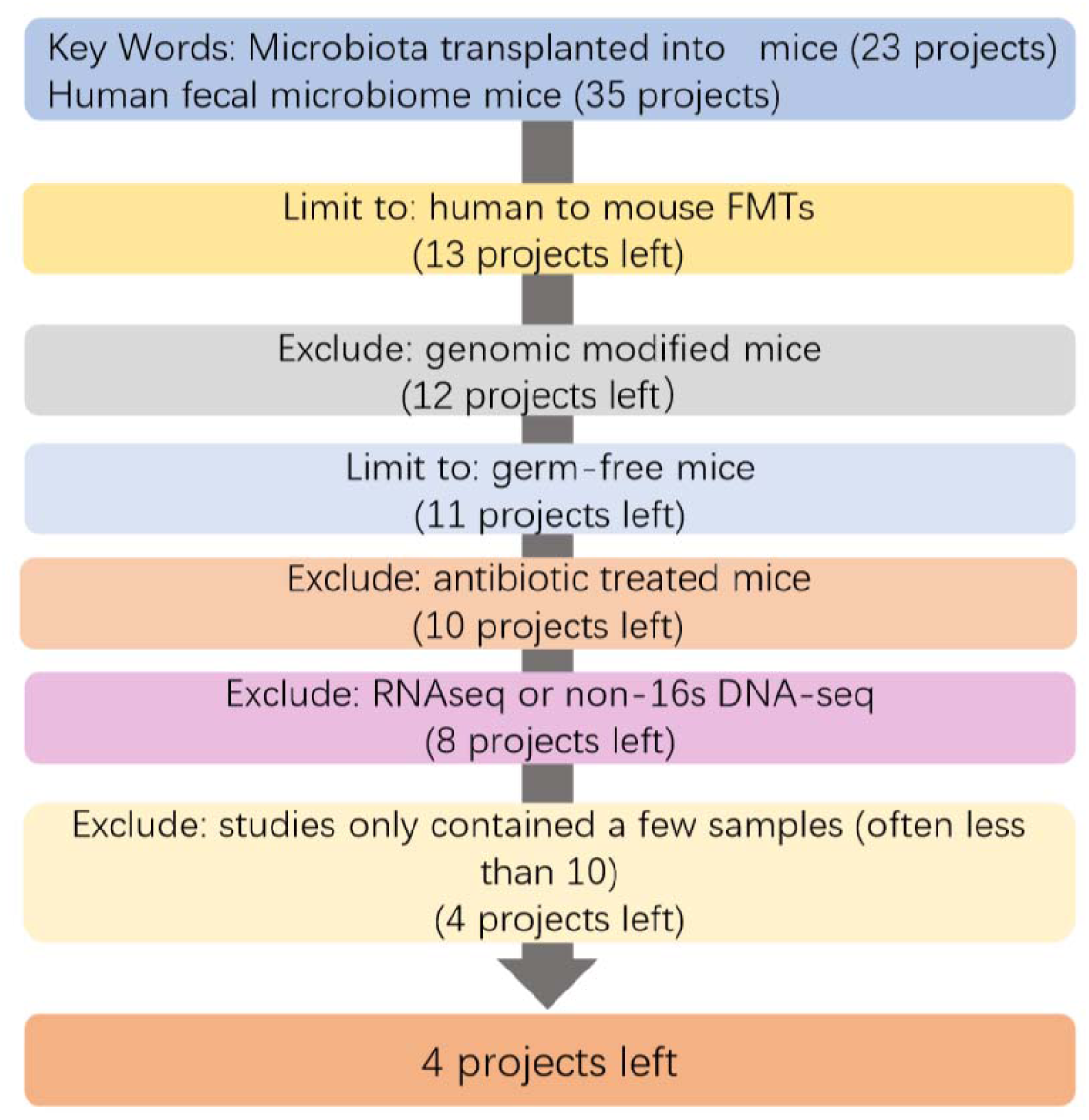
An extensive search of public data for human to mouse fecal transplantation experiments. A graphical workflow showed our searching and excluding criteria.

**Table 2.** a lists of studies (with their NCBI BioProject IDs) that met our search criteria.

| NCBI<br>BioProject<br>Accession | Nr.<br>Human-mouse<br>pairs | Published Date | Phenotypes<br>of<br>donors | Reference |
| --- | --- | --- | --- | --- |
| PRJNA314018 | 420 | 2-Mar-2016 | Obese and healthy | [29] |
| PRJEB7604 | 161 | 28-Oct-2014 | Obese and healthy | [41] |
| PRJEB11697 | 184 | 15-Mar-2016 | Healthy | [19] |
| PRJEB15481 | 948 | 21-Dec-2016 | Healthy | [26] |

### Data processing, taxonomic assignments

We used FastQC (downloaded from http://www.bioinformatics.babraham.ac.uk/projects/fastqc/; ver 0.11.8) to evaluate the overall quality of the downloaded data, followed by the use of Trimmomatic [42] to remove vector sequences and low-quality bases. We used the single-ended sequencing reads directly for subsequent analysis, and merged the pair-ended reads first using Casper [43]. We then used Qiime [44] to check and remove possible chimeras.

We used Mapseq [45] to analyze the obtained clean data and assign taxonomic classification information to the reads. It was previously shown that at the genus level Mapseq has higher accuracy than other poplar tools such as Qiime [44] and Mothur [46]. Mapseq is also advantageous in our study as compared with *de novo* clustering methods. The clean reads after the removal of low-quality and/or sequencing-primer sequences often have uneven ends, 16S sequences belonging to the same species/genus cannot be reliably clustered together. Consequently the retention rate, i.e. the proportion of human gut microbes that can be found in the recipient mice after FMT, will be significantly inflated. This is also part of the reason that EBI Metagenomics [47], a popular metagenomic database and analysis platform, has recently adopted Mapseq as the main tool for taxonomy assignment. We removed reads with a cutoff value at the genus level less than 0.4 (the combined score) as recommended by the authors of Mapseq [45]. We then calculated relative abundances calculated at the genus and species levels for each sample, with the totaling abundances values of 100% respectively.

Supplementary Table 2 listed all human-mouse pairs used in this study, along with their NCBI run IDs in the NCBI SRA database, corresponding enterotypes, experimental conditions and diet types.

### Statistical analysis

We loaded all processed data to R (version 3.5.1; downloaded from https://www.r-project.org; accessed in July 2018) for further analysis. To focus more on abundant species/genus and avoid false dramatic fold-changes in the calculation due to low-abundance species, we removed genera and species with abundance less than 0.1% from subsequent analysis. In order to make the results more accurate, we also removed genera and species supported by less than 5 reads.

We used a web-based tool (http://enterotype.embl.de/enterotypes.html) to determine the enterotype type for each sample by using the relative abundances of that sample as input.

## Abbreviations

FMT: Fecal microbiota transplant
GF mice: Germ-free mice
FC: Fold change
ASD: Autism Spectrum Disorder

## Declarations

### Acknowledgements

We are very grateful to Qianhui Zhu and Chengwei Zhang for their help in downloading 16S DNA sequencing data.

### Availability of data and materials

All runIDs from the four NCBI BioProjects are provided in Additional file 3: Table S2, they can be download from NCBI.

### Authors’ contributions

Wei-Hua Chen conceived the project. Yanze Li collected and analyzed the data. Wenming Cao contributed to the data download and analysis. Na L Gao helped with setting up the data analysis pipelines especially for the raw data processing. Wei-Hua Chen and Yanze Li wrote and revised the manuscript.

### Ethics approval and consent to participate

Not applicable.

### Consent for publication

Not applicable.

## Supplementary tables

**Supplementary Table 1.**
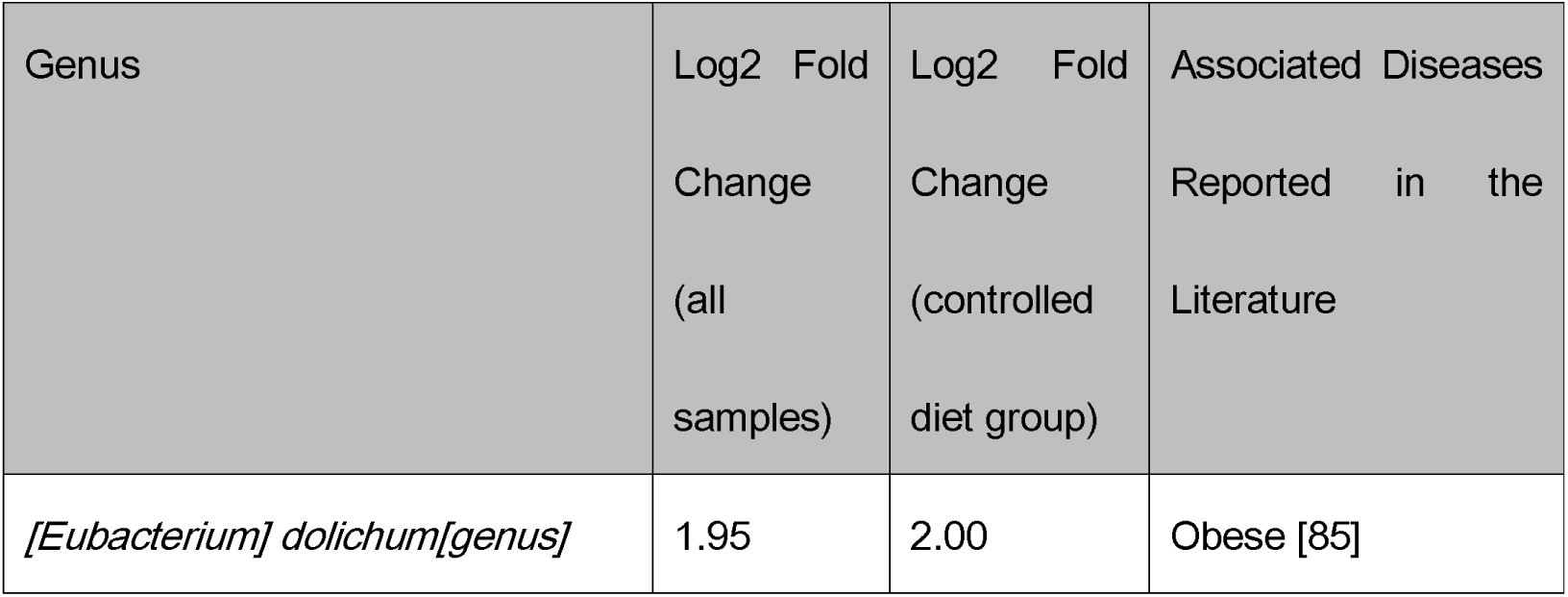
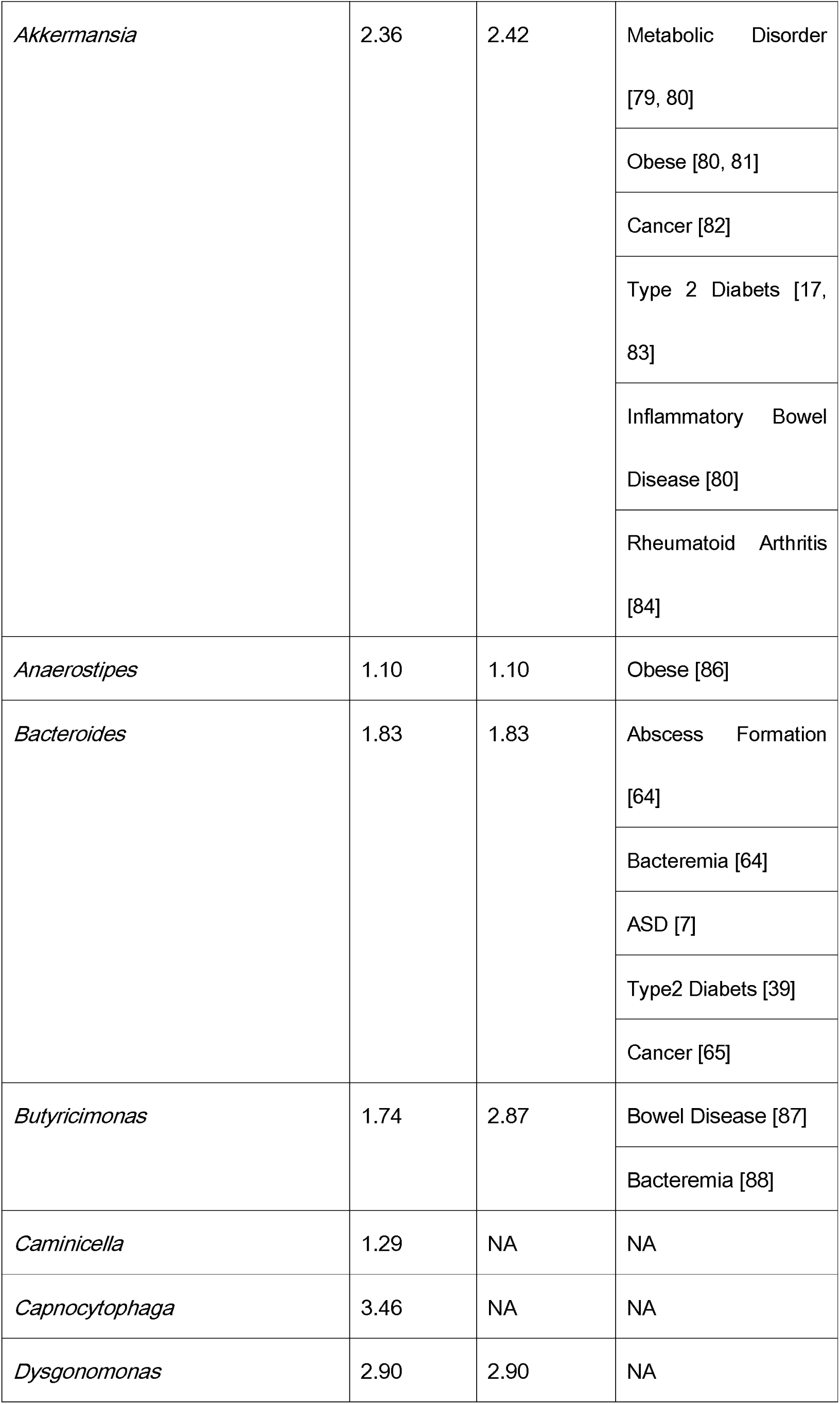
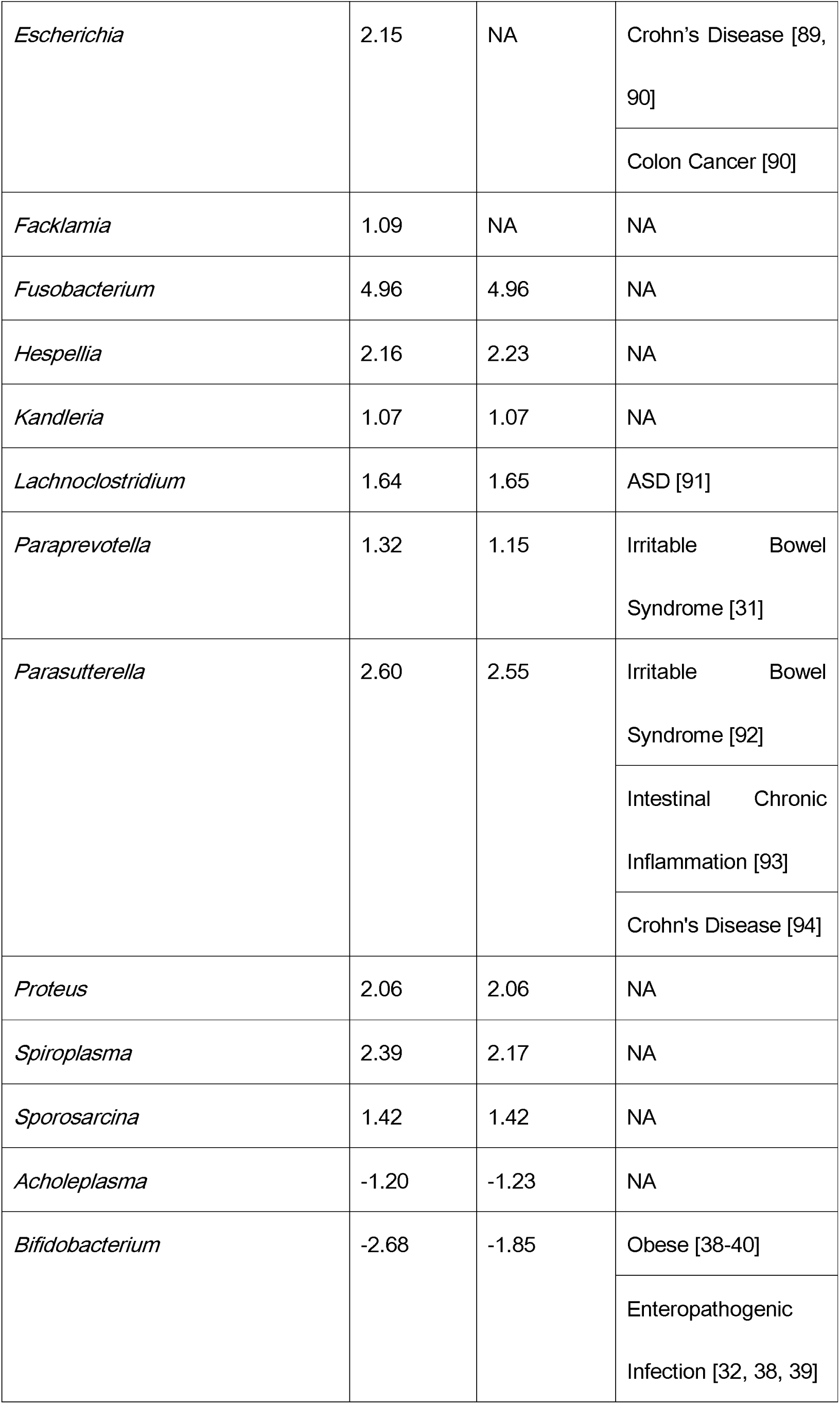
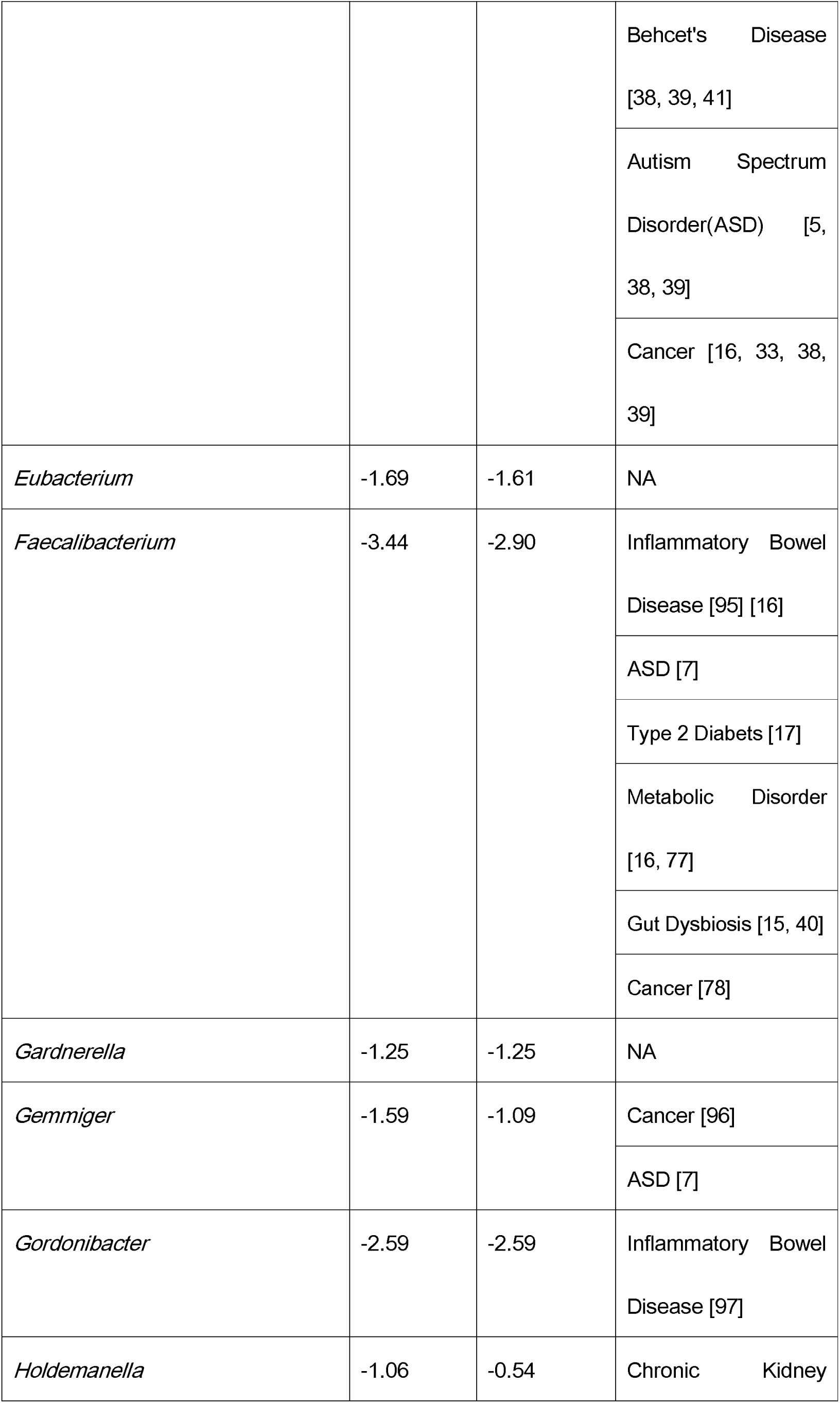
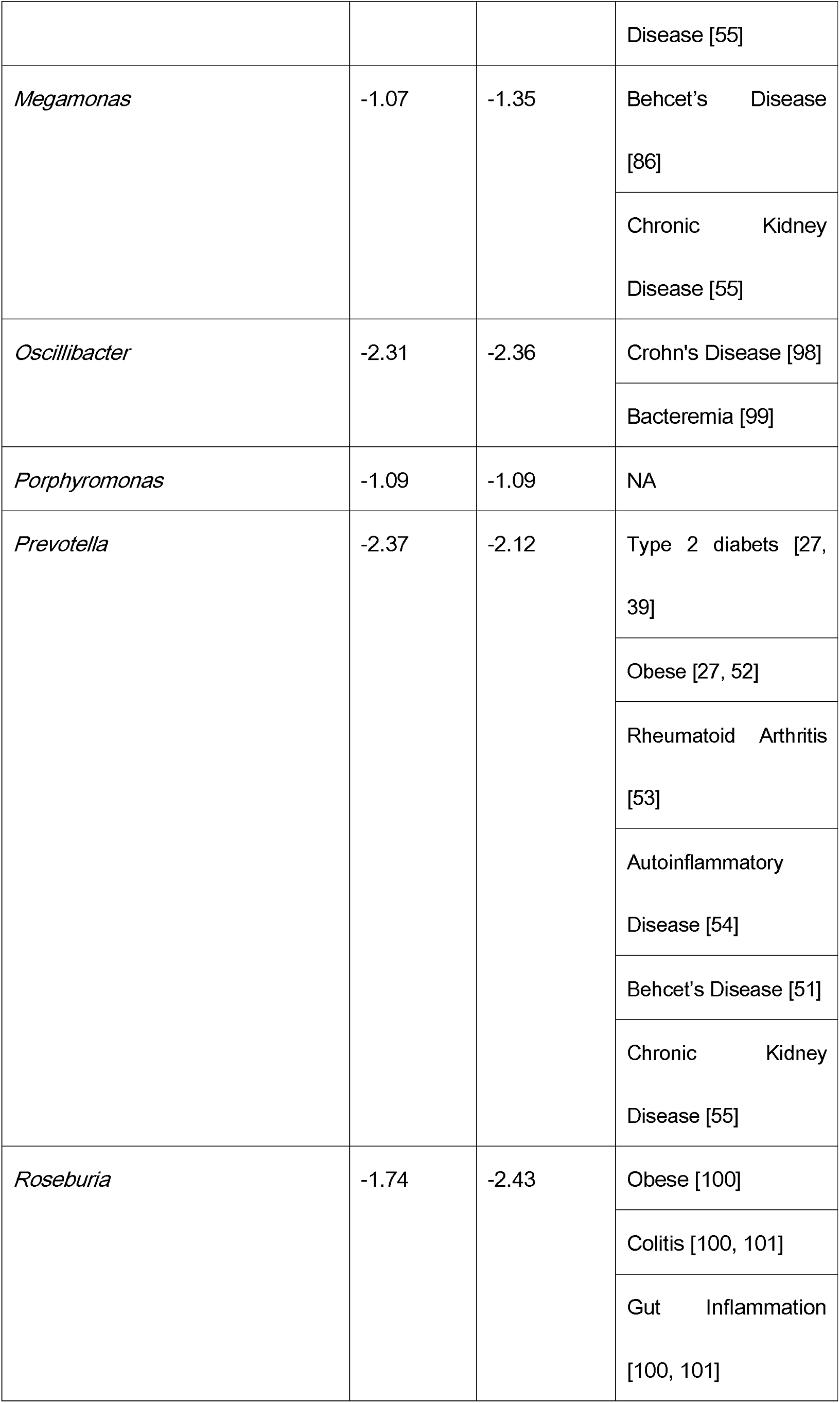
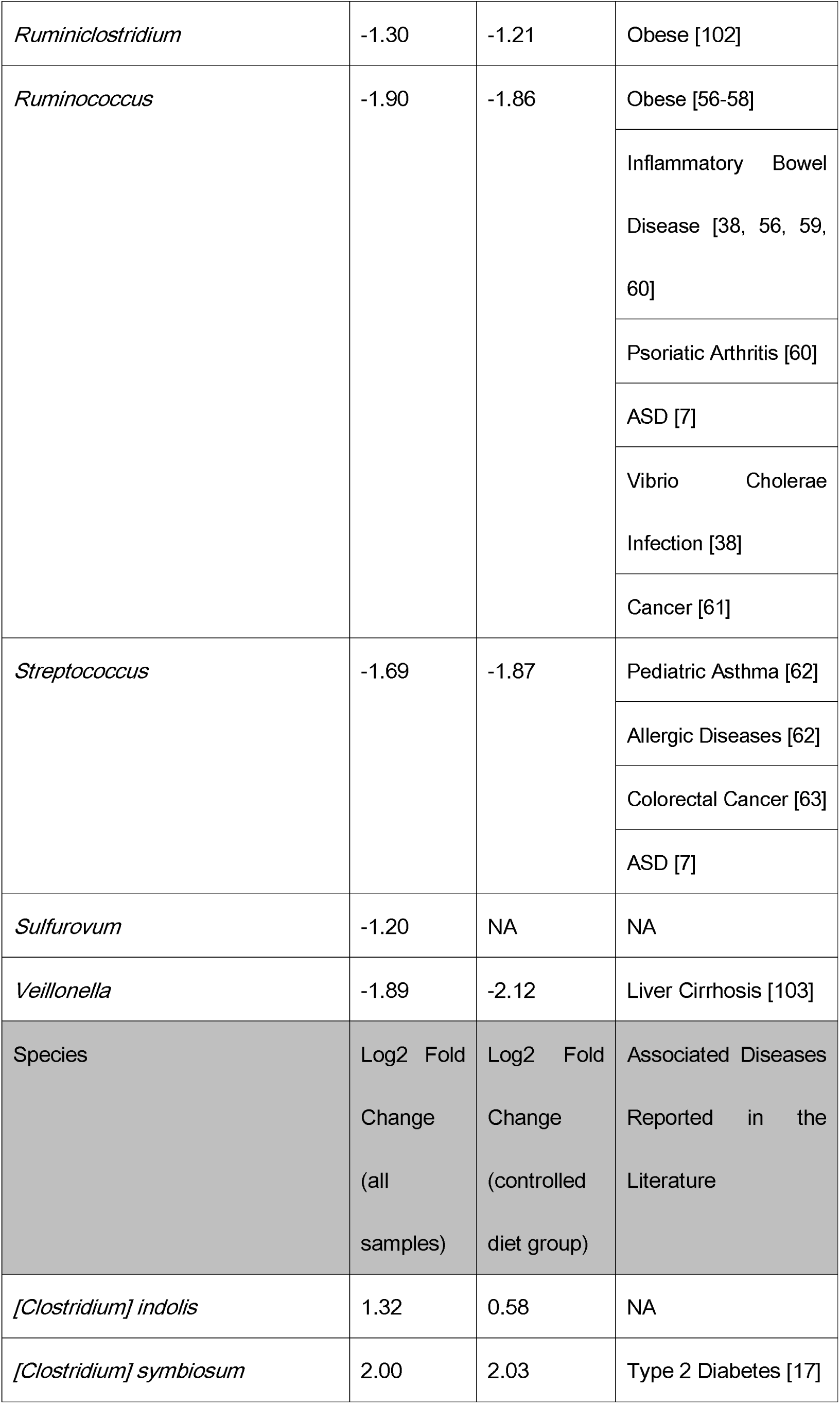
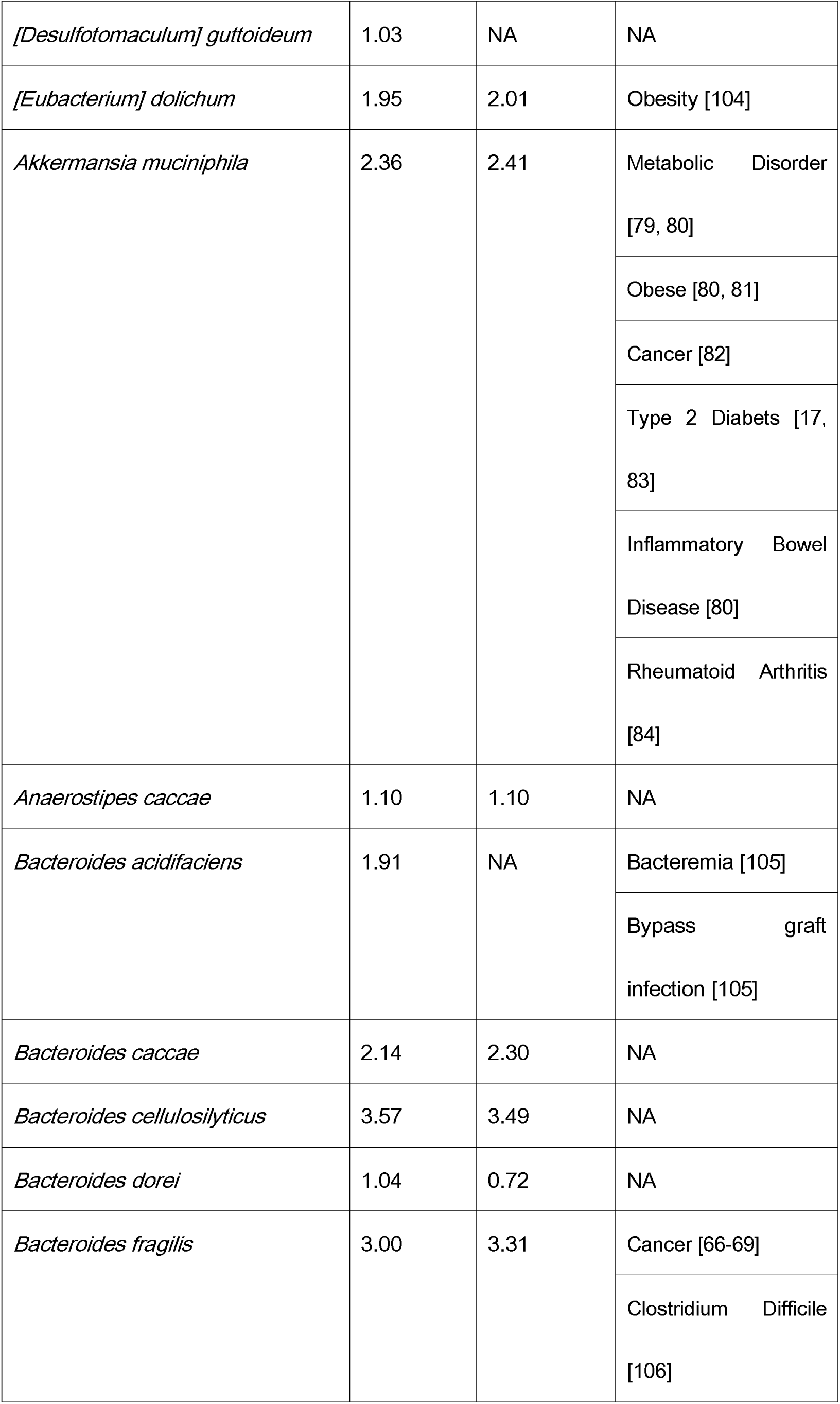
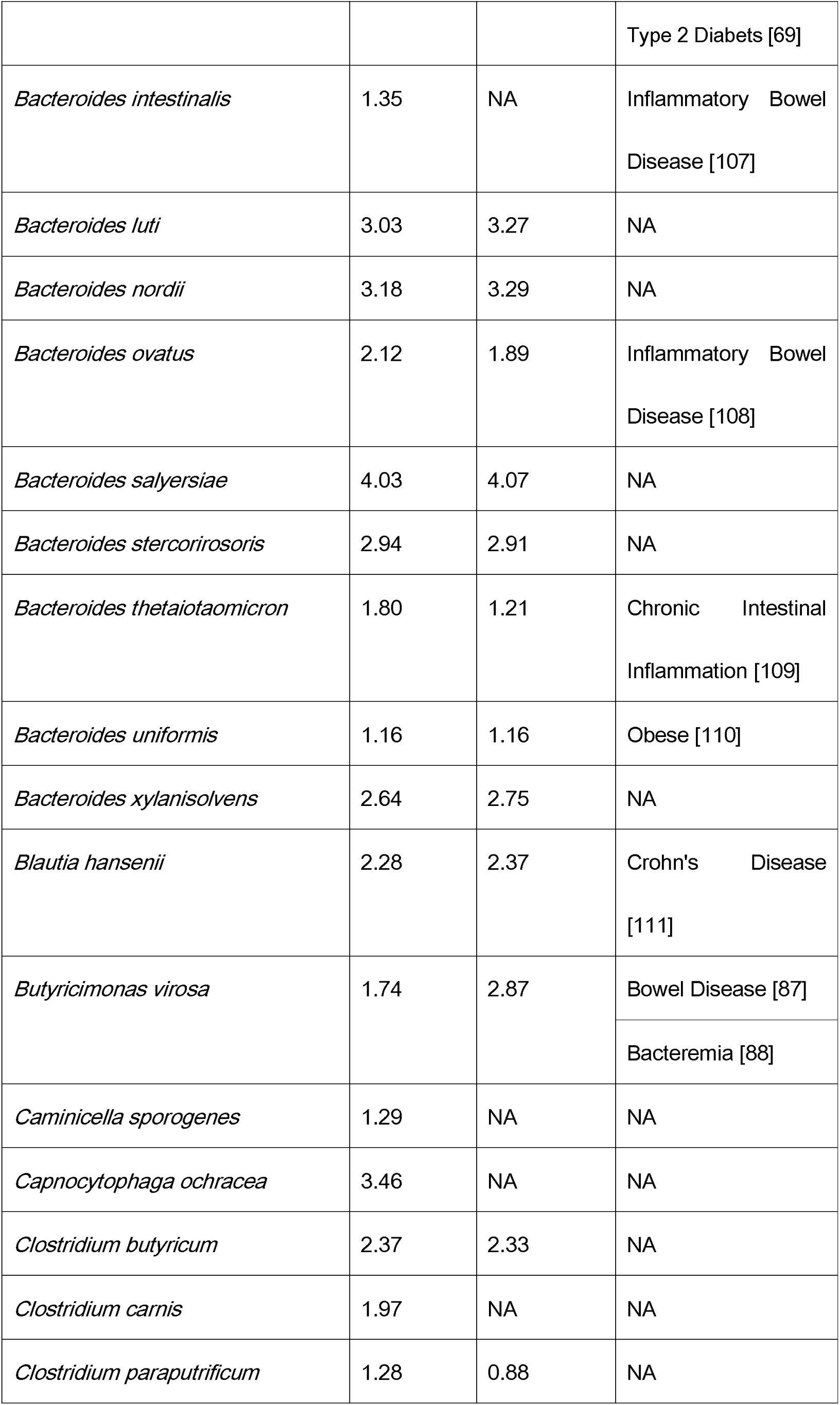
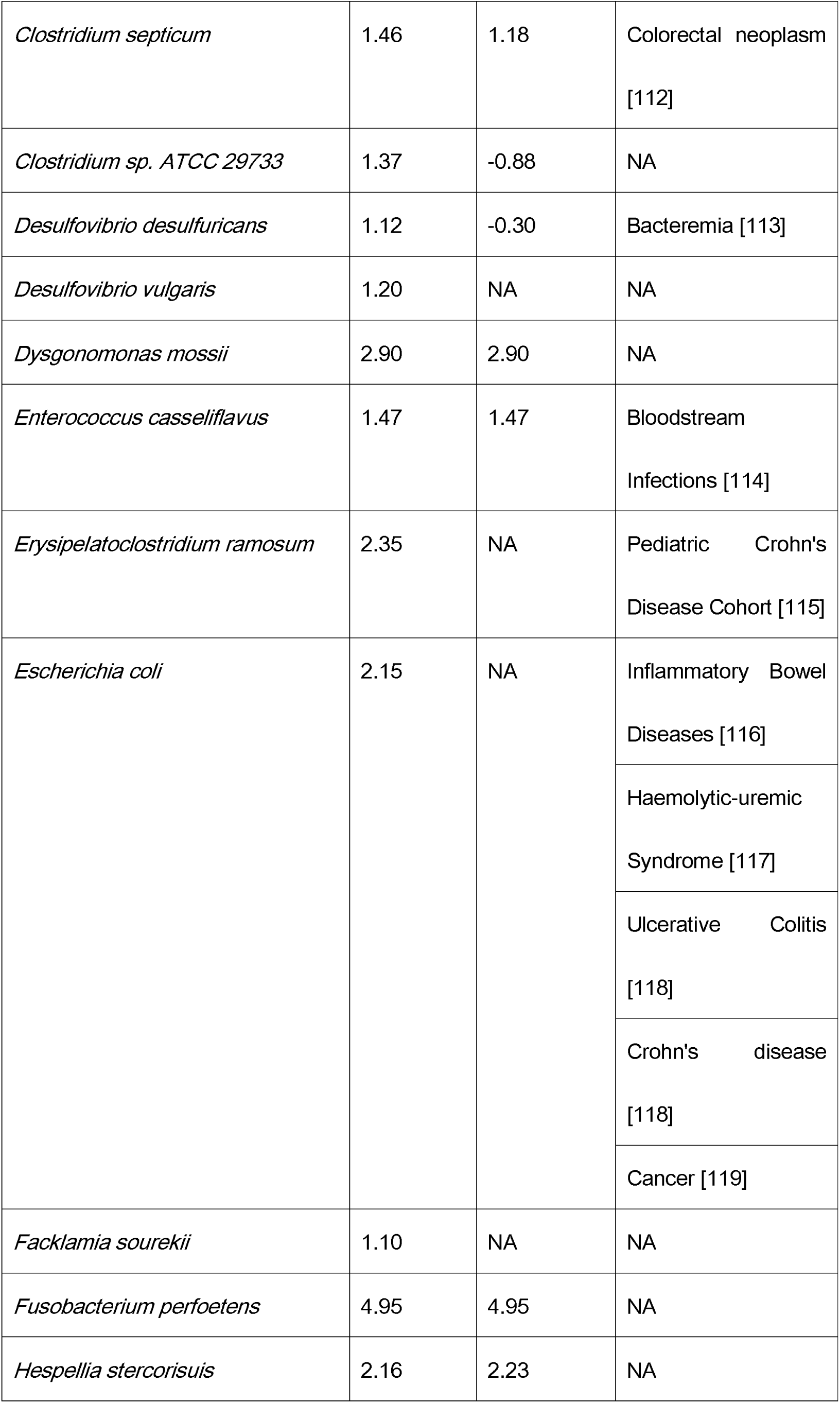
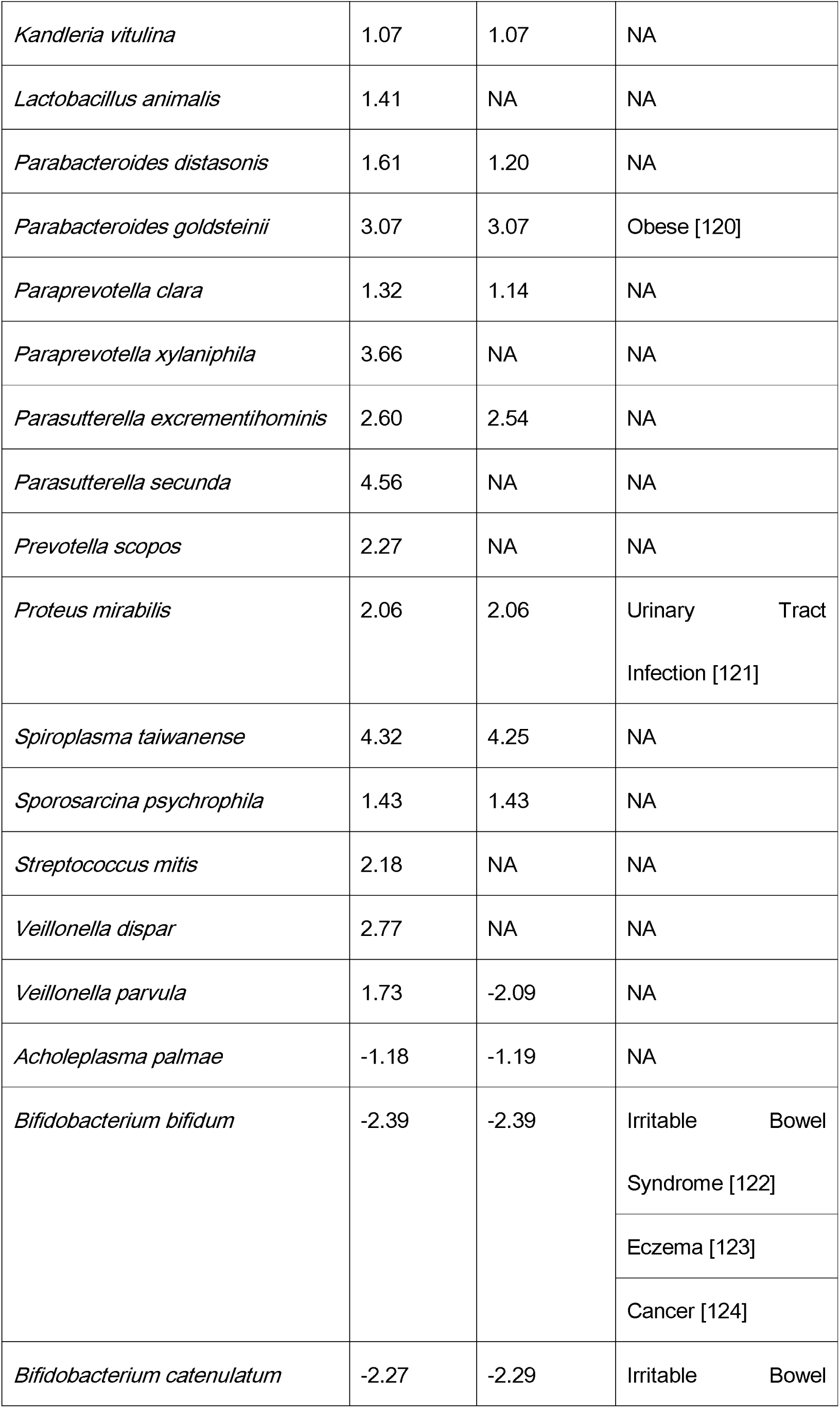
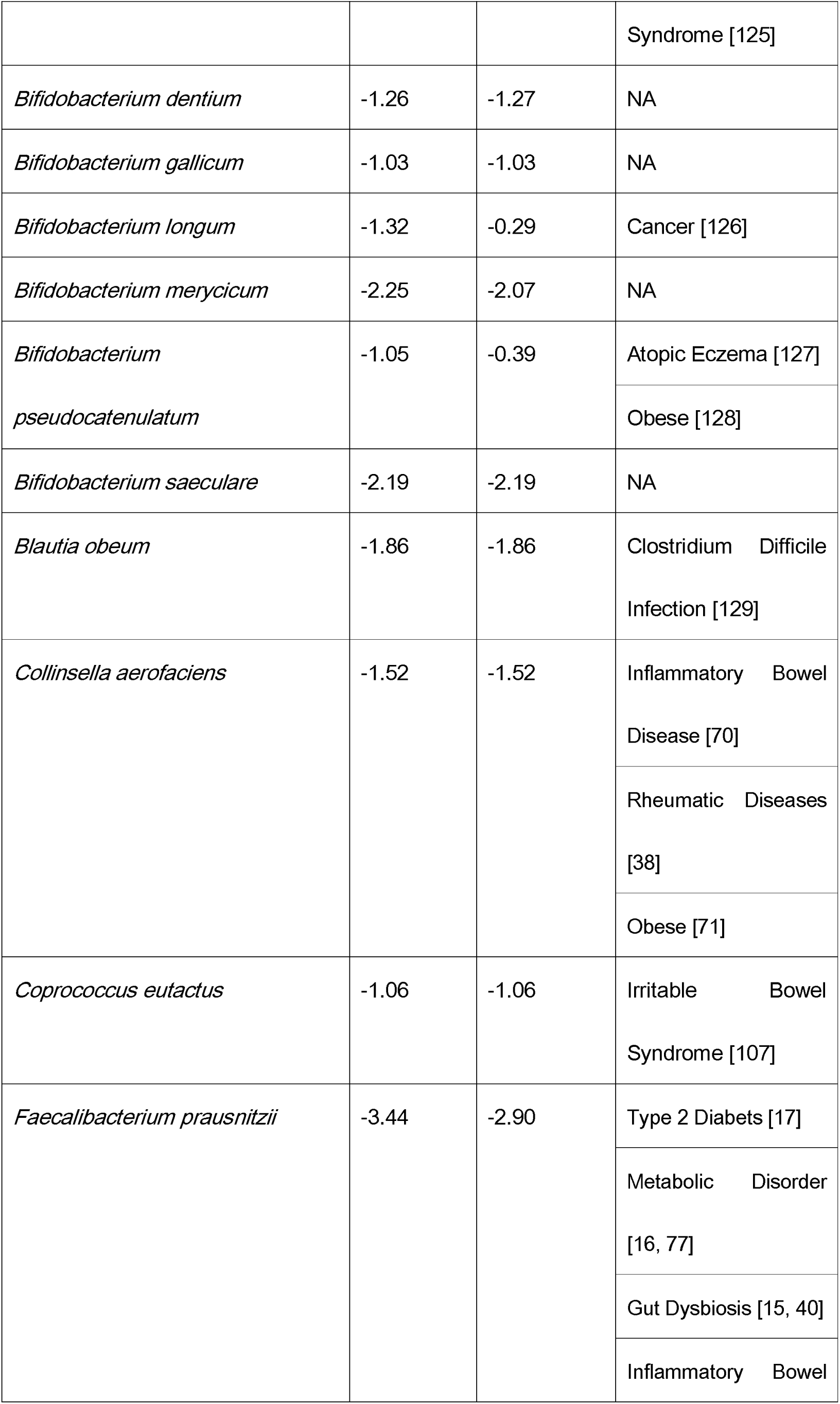
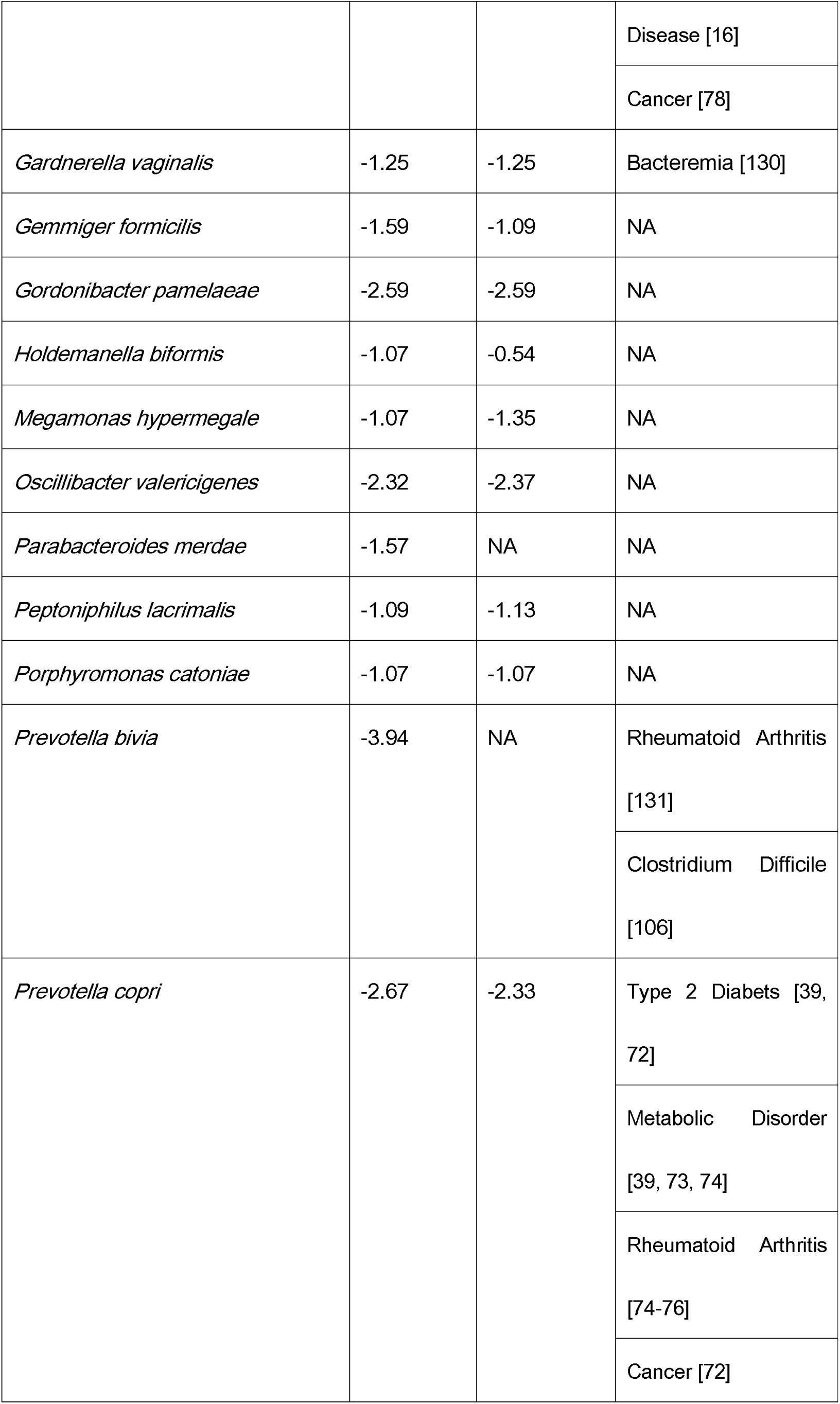
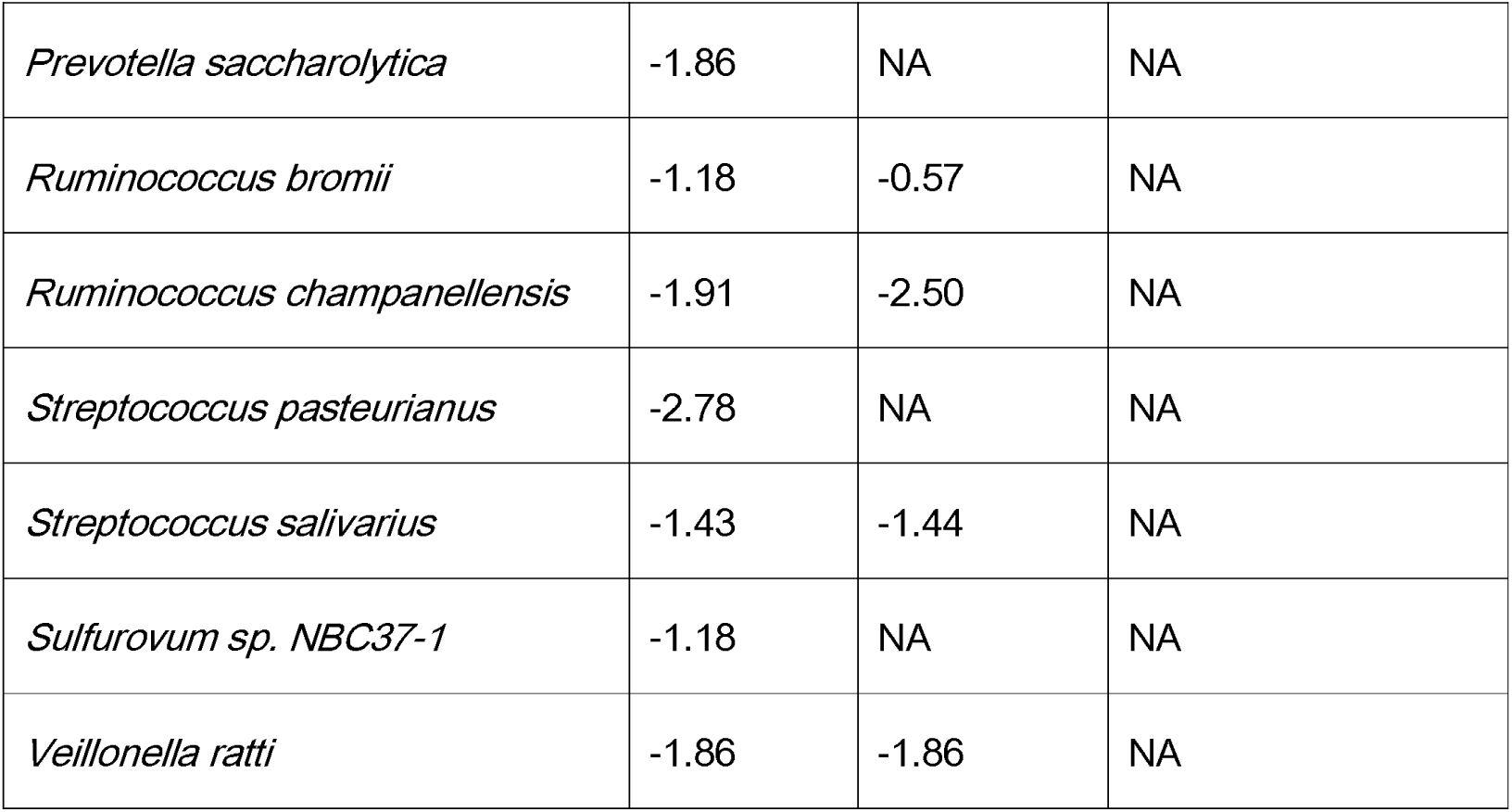
A complete list of variable taxa, their responses to diet changes and associations with human diseases.

**Supplementary Table 2 (submitted separately as an additional Excel file).** A complete list of human-mouse pairs (before and after FMT) and their corresponding experimental conditions, enterotypes and diet types

